# Tillage homogenizes soil bacterial communities in microaggregate fractions by facilitating dispersal

**DOI:** 10.1101/2023.03.08.531801

**Authors:** Jaimie R. West, Joseph G. Lauer, Thea Whitman

## Abstract

Soil aggregation physically protects soil organic matter and promotes soil carbon persistence through microaggregate formation and organo-mineral associations. Tillage is a ubiquitous disturbance to arable soil that disrupts aggregation, thus affecting microbial resource availability, soil microhabitat conditions, and microbial interactions. We investigated how tillage affects bacterial community composition of soil microaggregate fractions (53–250 µm), specifically the free microaggregate fraction in bulk soil, and the occluded microaggregate fraction from within macroaggregates, using two long-term tillage vs. no-tillage experiments in southern WI, U.S., that represent two different silt loam soils (Alfisol and Mollisol). We applied 16S rRNA gene amplicon sequencing to characterize the effects of tillage on microaggregate bacterial communities by relating compositional changes and ecological community assembly patterns to various tillage-driven changes in the soil environment, including aggregate size distribution and carbon content. Tillage homogenized soil bacterial communities, as quantified by increased compositional similarity at both within-plot and between-plot scales, and community assembly was increasingly influenced by homogenizing dispersal with tillage. We did not identify major distinctions between bacterial communities of the free and occluded microaggregate fractions, thus highlighting how soil microaggregates readily shift between these operationally defined fractions in temperate annual cropping systems, where the soil environment is subject to drastic seasonal changes that are exacerbated by tillage. With this study, we improve our understanding of the microbial response to soil disturbance, and thus the potential mechanisms through which disturbances like tillage affect soil carbon persistence.

**Highlights:**

- Tillage homogenized soil bacterial communities, within and between plots
- Homogenizing dispersal drove community assembly under tillage
- Free and occluded microaggregate fractions hosted similar communities

## 1. Introduction

Soil aggregation—the binding of soil particles and organic matter (Tisdall and Oades, 1982)—is a primary mechanism of soil organic matter (SOM) protection that promotes microbially-mediated soil carbon stability and persistence (Jastrow and Miller, 1998). Agricultural tillage, employed on over 60% of U.S. farmland (Zulauf and Brown 2019), disrupts aggregation, thus reducing physical protection of SOM and potentially increasing microbial mineralization of soil organic carbon (SOC) (Elliott, 1986; Paustian et al., 1997; Six et al., 1998; Schimel and Schaeffer, 2012). Improved understanding of the ecological factors that relate microbial community composition to changes in aggregation and SOC persistence under mixing disturbances will populate a key knowledge gap in soil microbial ecology (Wilpiszeski et al., 2019). These relationships may be pertinent in the highly protected microenvironments of soil microaggregates (< 250 µm in diameter), which are more stable and have lower turnover rates than macroaggregates (250–2000 µm in diameter) (DeGryze et al., 2006; Davinic et al., 2012; Totsche et al., 2018). Microaggregates are found both free in the bulk soil and occluded within macroaggregate structures (Oades, 1984; Totsche et al., 2018).

Microaggregates are cemented and glued together by physicochemical interactions and biomolecules that physically protect SOM, rendering it less accessible for microbial decomposition (Edwards and Bremner, 1967; Jastrow and Miller, 1998; Six et al., 2004; Totsche et al., 2018; Woolf and Lehmann, 2019). Microaggregate structures further inhibit microbial activity due to nutrient and oxygen limitation (Sexstone et al., 1985; Ranjard and Richaume, 2001). These same microhabitats that protect SOM from microbial decomposition are also disproportionately high in microbial abundance; an estimated 70% of soil bacteria live within microaggregates (Ranjard et al., 2000), despite the microaggregate fraction comprising perhaps 30–50% of arable soil by mass (Sheehy et al., 2015; Cates et al., 2016). Soil bacterial communities moderate the soil carbon cycle through decomposition of SOM and mineralization of SOC for food (Schimel and Schaeffer, 2012), while also contributing to persistent SOM through the production of microbial necromass, which has been shown to accumulate in soil (Simpson et al., 2004; Liang and Balser, 2008). Overall, the mechanisms that balance microbially mediated SOC persistence with carbon-consuming microbial activity in microaggregates (i.e., “microbial hotspots”, sensu Kuzyakov and Blagodatskaya, 2015) are not well-understood (Six et al., 2004; Wilpiszeski et al., 2019).

Agricultural tillage disrupts the fine roots and fungal hyphae that stabilize macroaggregates (Tisdall and Oades, 1982; Elliott, 1986; Six et al., 1998), thus decreasing mean aggregate size and/or proportion of aggregated soil (Frey et al., 1999; Six et al., 1999; Al-Kaisi et al., 2014; Zheng et al., 2018). Because microaggregates can form within the protective environment of a macroaggregate (Oades, 1984; Angers et al., 1997), tillage reduces the potential for occluded microaggregate development within macroaggregates and, critically, the rate at which SOM is potentially stabilized in microaggregates (Six et al., 2000a). Further, as macroaggregates destabilize, their occluded microaggregates become more freely connected to the bulk soil environment, increasing resource diffusion (e.g., oxygen and extracellular enzymes) and decomposer pressure (Six et al., 1999; Garland et al., 2018; Piazza et al., 2020). Through these mechanisms, tillage has been associated with decreases in total SOM content (Elliott, 1986), SOM residence time (Paustian et al., 2000), SOC content (Paustian et al., 1997; Al-Kaisi et al., 2014; Zheng et al., 2018), aggregate-occluded particulate organic matter (POM) (Six et al., 1999), microbial biomass (Zuber and Villamil, 2016), and microbial necromass accumulation (Simpson et al., 2004). Though these effects are well-documented, they are typically only noted in the top 5 or 10 cm of soil (Frey et al., 1999; Six et al., 1999; Simpson et al., 2004; Zheng et al., 2018), and some work has suggested that tillage does not decrease total SOC across the full soil profile (Powlson et al., 2014; Ogle et al., 2019). There is also evidence that minimum tillage practices can be equally beneficial as no-tillage regarding SOC and microbial necromass accumulation, by incorporating nutrients and alleviating compaction (Sae-Tun et al., 2022).

A recent meta-analysis demonstrated that the occluded microaggregate fraction preferentially accumulates SOC at a higher rate than the free microaggregate and other soil fractions (King et al., 2019). One study found over 90% of the increase in SOC content in no-tillage as compared to conventional tillage systems was attributable to the occluded microaggregate fraction, across soils of various clay mineralogies (Denef et al., 2004), while another study found that the occluded microaggregate fraction contributed 49–112% of the increase in SOC following a shift to no-tillage across the U.S. (Six and Paustian, 2014). Together, these results indicate a higher capacity for SOC persistence in the occluded versus the free microaggregate fraction.

The effects of tillage on soil microenvironments (e.g., aggregate size and porosity), and the resulting redistribution of resources (e.g., oxygen, water, biomass), suggests that tillage also alters soil microbial community composition and function (Bhattacharyya et al., 2021). Tillage- driven decreases in aggregate size may select for more oligotrophic communities due to lower substrate and oxygen availability (Trivedi et al., 2017), though some have found fast growing, copiotrophic competitors to dominate soil communities under tillage or disturbance (Srour et al., 2020; West and Whitman, 2022). Tillage also shifts communities towards bacterial dominance and away from fungal dominance, which is thought to contribute to SOC loss (Six et al., 2006). Understanding changes to microbial communities under a given management practice, such as tillage, is essential for improving predictions of SOC persistence and storage.

In the limited number of studies that have applied high-throughput sequencing to aggregate fractions, distinct and more diverse bacterial communities are supported by the free microaggregate fraction than the macroaggregate fraction (Trivedi et al., 2017; Bach et al., 2018; Upton et al., 2019). One study that separately assayed free and occluded microaggregate fractions found similar community compositions in the free and occluded microaggregate fractions (53–250 µm), yet suggested that copiotrophic bacteria live in association with free microaggregates whereas oligotrophic bacteria are characteristic of occluded microaggregates (Biesgen et al., 2020). This assessment is consistent with the idea that free microaggregates have higher resource availability, notably C and oxygen, that may support copiotrophic microorganisms, whereas occluded microaggregates may be more insulated from perturbation, resource fluxes, and decomposers, as evidenced by increased SOC persistence (King et al., 2019). Another study found that microbial necromass concentration was greater in the occluded microaggregate fraction under no-tillage as compared to tillage, whereas there was no difference in the free microaggregate fraction under tillage versus no-tillage (Simpson et al., 2004). These results suggest that tillage-related impacts on macroaggregate formation and turnover (and, thus, occluded microaggregates) extend to microbial community composition (Six et al., 2004).

In this study, we assess differences in community composition of the free and occluded microaggregate fractions to tease out the ecological processes governing community assembly, in order to better predict how management-driven changes to soil structure affect soil communities. The community assembly processes (Vellend, 2010) of interest are as follows: Dispersal describes the generally stochastic movement and establishment of organisms in space, and may occur in soil via physical disturbance or mass flow of pore water (Zhou and Ning, 2017). Homogenizing dispersal increases compositional similarity between communities, whereas dispersal limitation increases compositional differences between communities, and may allow for stochastic demographic changes to community composition — termed ‘drift’ (Stegen et al., 2013). Selection refers to deterministic or niche-based processes dictated by biotic factors, such as inter-taxa fitness differences, and abiotic factors, such as environmental filters (Hutchinson, 1957). Homogeneous selection decreases phylogenetic differences between communities due to community assembly under similar conditions or filters (Dini-Andreote et al., 2015). Variable selection increases phylogenetic differences between communities due to variable conditions (Stegen et al., 2015). To statistically infer the relative influences of these community assembly processes in soil microbial communities, Stegen et al. (2012, 2013, 2015) developed a null modeling approach that compares observed phylogenetic distances and dissimilarity metrics between communities to null models of stochastically assembled communities. A more recent approach separately assesses community assembly processes within phylogenetically related ‘bins’ of OTUs, thus enabling representation of various assembly processes that may influence subsets of community members (Ning et al., 2020). These approaches have not yet been used to directly compare the effects of tillage on community assembly, let alone at the microaggregate fraction scale.

We sought to better understand how bacterial communities are affected by tillage, as modulated through soil aggregation. We collected soil samples in no-tillage and chisel-plowed tillage plots from two long-term studies in southern Wisconsin, U.S., and related soil properties to tillage- driven differences in bacterial community composition, diversity, and community assembly processes of the bulk soil, free microaggregate, and occluded microaggregate fractions, using 16S rRNA gene amplicon sequencing. In addition to expecting standard responses to tillage including decreased SOC and aggregation, we hypothesized that the free microaggregate and occluded microaggregate fractions would support distinct bacterial communities, and demonstrate differences due to tillage treatments. Specific hypotheses included: H1) With tillage, community assembly would be driven by homogenizing dispersal and homogeneous selection, whereas in the no-tillage system, community assembly would be driven by dispersal limitation and variable selection. H2) The occluded microaggregate fraction would demonstrate stronger evidence for dispersal limitation, whereas the free microaggregate fraction would demonstrate stronger evidence for homogeneous selection. H3) Tillage would increase sample-to-sample similarity in community composition (i.e., lower beta diversity). Better understanding microbial community composition and assembly in microaggregate environments will improve efforts to understand mechanisms of SOC persistence, thus contributing to climate change resilience (Paustian et al., 2000), ecosystem services, and crop productivity (Janzen, 2006).

## 2. Methods

### 2.1 Soil collection

Soil was sampled from two separate long-term tillage studies located at 1) the University of Wisconsin (UW) Arlington Agricultural Research Station in Arlington, WI, U.S., (43°17’56”N, 89°21’11”W, 314 m a.s.l.) on a Plano silt loam soil (fine-silty, mixed, superactive, mesic Typic Argiudoll), under a corn (*Zea mays* L.) – soybean (*Glycine max* L.) rotation; and, 2) the UW Lancaster Agricultural Research Station in Lancaster, WI, U.S., (42°49’53”N, 90°47’35”W, 313 m a.s.l.) on a Fayette silt loam soil (Fine-silty, mixed, superactive, mesic Typic Hapludalfs), under a continuous corn rotation. The tillage study at Arlington, WI was established in 1987 with a no-tillage treatment, in which crops are planted directly into the undisturbed residue of the previous crop, and a tillage treatment, which consists of fall chisel plow followed by two spring field cultivator passes prior to planting. Further details regarding management practices and agronomic findings have been reported (Pedersen and Lauer, 2003; Chamberlain et al., 2021).

The tillage plots at Lancaster, WI were established in 1993, consisting of no-tillage and tillage treatments, the latter of which consists of fall chisel plow and a spring field cultivator pass prior to corn planting. The Lancaster plots have been used for various research projects over the years (e.g., Gupta et al., 2004; Dolliver and Gupta, 2008), which sometimes included manure application treatments (1993–1997, 2003–2005, 2014) or corn fungicide treatments (2008–2010).

Soil was sampled once in each location (23 October 2021 at Arlington, WI and 6 November 2021 at Lancaster, WI), following corn grain harvest and prior to fall tillage. At Arlington, three plots were sampled for each treatment, collecting five intact cores per plot for a total of 15 cores per treatment. Due to our interest in discerning dispersal processes, our sampling design focused on ensuring relatively high spatial proximity of individual cores within a given plot. Soil cores were 7.9 cm dia, evenly spaced just within the perimeter of a 48 cm dia circle; distance between adjacent cores was approximately 15 cm. As detailed below, the top 5 cm was analyzed to target soil under the greatest intensity of tillage disturbance. Lancaster was sampled in the same fashion, but only two plots per treatment were used for analysis (see section 2.7), for a total of ten cores per treatment. Intact cores were temporarily kept in a cooler, and then held at 4 °C for up to ten days until sample processing.

### 2.2 Aggregate size fractionation and sample processing

To assess variability in community composition and community assembly at a relatively small spatial scale, each core was processed separately. The top 5 cm of each field-moist soil core were gently passed through a 2 mm sieve (henceforth referred to as “bulk” soil). Then, 80 g of this field-moist bulk soil was subjected to aggregate size fractionation via wet sieving (Elliott, 1986) to isolate the macroaggregate fraction (250 µm–2000 µm), free microaggregate fraction (53 µm– 250 µm), and the silt + clay-sized fraction (< 53 µm). Then, 20 g of the moist macroaggregate fraction was separated into occluded fractions via rapid shaking with glass beads to break up the macroaggregates under water flow to encourage liberated microaggregates and other material to pass through a 250 µm sieve, as previously described (Six et al., 2000a, 2002); macroaggregate-occluded fractions included the occluded microaggregate fraction (53 µm–250 µm), occluded silt + clay-sized fraction (< 53 µm), and occluded coarse POM + coarse sand-sized fraction (250 µm – 2000 µm). Other modifications to the wet sieving method included a slaking for two minutes prior to the first wet sieving step (Arlington samples only) and draining each wet sieved fraction for two minutes prior to subsampling as described below. The largely unaggregated Lancaster soil samples did not undergo slaking, and required wet sieving of an additional 80 g of bulk soil to obtain enough macroaggregate soil for the occluded fraction separation step. The primary objective of fractionation was to isolate the free and occluded microaggregate fractions, but the relative dry mass of each size fraction was also determined. Correction for sand content of aggregate fractions was not performed and thus all size fractions also include the primary mineral particles of that size.

Moisture content was estimated for bulk soil, the wet-sieved macroaggregate fraction, the free microaggregate fraction, and the occluded microaggregate fraction by drying subsamples in a 60 °C oven for 24 hours. Bulk, free microaggregate, and occluded microaggregate soil was subsampled for DNA extraction (see section 2.4), and bulk soil was also subsampled to measure soil respiration (see section 2.3). The remaining wet-sieved soil was washed from each sieve (or washbasin) into aluminum pans to determine the dry mass of each fraction. Overall recovery (macroaggregate + free microaggregate + [silt + clay] fractions) was 99% for both treatments at both sites, and macroaggregate recovery (occluded microaggregate + [occluded silt + clay] + occluded coarse POM) was 101% for tillage treatments, and 96–97% for no-tillage treatments.

### 2.3 Soil analysis

The bulk soil (sieved to < 2 mm), macroaggregate, free microaggregate, and occluded microaggregate fraction subsamples that were retained for dry mass conversion were ground to a powder and used to quantify total soil carbon and nitrogen by flash combustion with a Flash EA 1112 CHN Automatic Elemental Analyzer (Thermo Finnigan, Milan, Italy) and soil pH (excluding the macroaggregate fraction). Soil pH was determined from the supernatant of a 1:1 soil:CaCl_2_ (0.1 M) slurry using a micro pH electrode (modified from Braus and Whitman, 2021). For routine soil analysis, a composite soil sample representing each treatment was comprised of an equal mass of bulk soil from each plot. Samples were submitted to the UW Soil and Forage Analysis Lab (Madison, WI, U.S.) to determine soil texture, organic matter content, pH, and plant-available P, K, Ca, and Mg. Routine soil properties can be found in Table S1.

Soil respiration (CO_2_ evolution) from fresh sieved soil was measured using the MicroResp system (James Hutton Ltd., Aberdeen, Scotland), following general instructions for use (Campbell et al., 2003) without added substrate. At the time of aggregate fractionation, 300 mg of freshly sieved (< 2 mm), field-moist soil from each soil core was placed into each of six wells of a deep-well plate. Wells were covered and stored at 4 °C for up to six hours. Each deep-well plate, containing soil from up to ten different cores, was covered in parafilm and firmly tapped on the benchtop 20 times to repack soil and minimize large air pockets. The deep-well plate was then incubated at 25 °C in a CO_2_-free environment overnight (approximately 16 hours), to help deplete CO_2_ from the well headspace and soil air. The following day, a colorimetric detection plate was read at absorbance wavelength 570 nm using a BioTek Synergy 2 spectrophotometer microplate reader (time = 0). After confirming that all wells had highly similar readings (< 5 % coefficient of variance), the detector plate was inverted over the deep-well plate, connected by the 96-well seal, and clamped together. After six hours of incubation at 25 °C, the clamp set-up was dismantled, and the colorimetric plate was immediately re-read to determine CO_2_ evolution. The colorimetric agar in the detection plates was made per the MicroResp manual instructions (version 4), and CO_2_ evolution was calculated following the manual’s instructions, with a minor modification to subtract CO_2_ values for a set of empty deep wells in each plate (blanks).

### 2.4 DNA extraction and 16S rRNA gene sequencing

Total genomic DNA was extracted from bulk soil, free microaggregate, and occluded microaggregate soil fractions using the DNeasy PowerLyzer PowerSoil Kit (Catalog No. 12855, Qiagen, Germantown, MD), following manufacturer’s instructions. We used 250 mg samples of field-moist bulk soil for DNA extraction, but, due to the wetness of the microaggregate fractions following wet sieving, we used 450 mg samples of these fractions to capture the same dry-mass equivalent of 250 mg of field-moist bulk soil, based on preliminary testing. The microaggregate samples were transferred directly from the drained sieves into the DNA extraction tubes, which were immediately frozen at −20 °C, and stored at −80 °C for up to three months prior to DNA extraction. Complete library preparation details can be found in the Supplementary Information. Briefly, the 16S rRNA genes of extracted DNA were amplified in triplicate using PCR. Variable region V4 of the 16S rRNA gene was targeted using forward primer 515f and reverse primer 806r (Walters et al., 2016). Primers also contained barcodes and Illumina sequencing adapters (Kozich et al., 2013). The following reagents comprised each 25 μL PCR reaction: 12.5 μL Q5 Hot Start High-Fidelity 2X Master mix (Catalog No. M0494, New England BioLabs, Ipswich, MA), 1.25 μL 515f forward primer (10 mM), 1.25 μL 806r reverse primer (10 mM), 1 μL DNA extract, 1.25 μL Bovine Serum Albumin (20 mg/mL; Catalog No. 97064-342, VWR International, Radnor, PA), and 7.75 μL PCR-grade water. The plate was sealed and briefly centrifuged prior to 30 PCR cycles on an Eppendorf Mastercycler nexus gradient thermal cycler (Hamburg, Germany) using the following parameters: 98 ^∘^C for 2 min + 30 × (98 ^∘^C for 10 seconds + 58 ^∘^C for 15 seconds + 72 ^∘^C for 10 seconds) + 72 ^∘^C for 2 min and 4 ^∘^C hold.

Amplified DNA was confirmed via gel electrophoresis, then normalized and purified, prior to paired-end 250 base pair sequencing on an Illumina MiSeq sequencer at the UW–Madison Biotech Center. To obtain high coverage, the same library was sequenced twice under identical conditions, and total reads were pooled for each sample after processing as described next.

Sequencing data were processed using a QIIME2 (Bolyen et al., 2019) pipeline, with DADA2 (Callahan et al., 2016) as the operational taxonomic unit (OTU, or amplicon sequence variant)- picking algorithm, and taxonomy assignment using the SILVA 132 reference database (Quast et al., 2013; Yilmaz et al., 2013). This yielded 10,102,355 demultiplexed sequences, which was reduced to 6,307,452 after denoising, with a mean length of 227 base pairs (SD = 2.2). Excluding extraction blanks, a total of 18,180 OTUs were identified. Amplicon sequences are available in the National Center for Biotechnology Information (NCBI) Sequence Read Archive (SRA) under accession [TBD]. Our primers targeted both bacteria and archaea, but because our communities were dominated by bacteria (94.5% of total reads), for simplicity, we will refer to bacteria when discussing communities in this manuscript. Over 99% of archaeal reads represented the phylum *Crenarchaeota*.

### 2.5 Data analysis

Data analysis was performed in R (R-Core-Team, 2018), using *ggplot2* (Wickham, 2016) for data visualization. The R code used to perform these analyses and to create the following figures is available at https://github.com/jaimiewest/Soil-Disturbance-Tillage. To test for a significant effect of tillage treatment on proportion of soil in each fraction, C content of each fraction, and respiration, we used ANOVA followed by Tukey’s HSD *post-hoc* comparison for significant results (*p* < 0.05). To test for a significant effect of tillage treatment, soil fraction, or interaction of these factors on soil C content, soil N content, soil C:N ratio, and soil pH, we performed ANOVA as described above. Unless otherwise noted, reported *p* values refer to ANOVA tests.

Community composition was visualized using principal coordinates analysis (PCoA) created with the *ordinate* function in the *phyloseq* package for R (*phyloseq::ordinate*) (McMurdie and Holmes, 2013) using Bray-Curtis dissimilarities (Bray and Curtis, 1957) of Hellinger-transformed relative abundance data (Legendre and Gallagher, 2001). To test for a significant effect of tillage treatment, soil fraction, or interaction of these factors on community composition, we used permutational multivariate analysis of variance (PERMANOVA) to partition Bray-Curtis dissimilarity matrices among sources of variation using *vegan::adonis2* (Anderson, 2001). A significant result (*p* < 0.05) was subjected to *post-hoc* pairwise factor comparisons, adjusting p-values using the Benjamini-Hochberg method (Benjamini and Hochberg, 1995) to identify significant differences. To compare differences in community composition due to tillage treatment or soil fractions, we tested for homogeneity of multivariate dispersions (PERMDISP; *vegan::betadisper*) (Anderson, 2006), using ANOVA to test the distances to group spatial median. Further, we also evaluated the effect of tillage treatment on dispersion of free and occluded microaggregate fraction communities within each soil core. To describe richness, we used the weighted linear regression model of OTU richness estimates, which weights observations based on variance, using *breakaway::betta* (Willis et al., 2017). We also calculated Faith’s phylogenetic diversity (PD) (Faith, 1992; Pérez-Valera et al., 2015) to assess differences in phylogenetic distance (i.e., sample branch length) using *picante::pd* (Kembel et al., 2010).

To further understand changes in community composition, we calculated the weighted mean predicted 16S rRNA gene copy number (Nemergut et al., 2016), which has been shown to correlate with potential growth rate (Klappenbach et al., 2000) and disturbance (Whitman et al., 2019; West and Whitman, 2022), and compared tillage treatments and soil fractions using ANOVA and *post-hoc* testing as described above. 16S rRNA gene copy numbers were predicted using the ribosomal RNA operon database (rrnDB) (Stoddard et al., 2015).

After evaluating our key questions, we used differential abundance to identify significant treatment-driven shifts in relative abundances of individual taxa as well as phyla. For this analysis, we compared the tillage treatments to each other (excluding taxa with mean relative abundance < 0.00001) and subjected those data to a beta-binomial regression model and “Wald” hypothesis test in *corncob∷differentialTest* (Martin et al., 2021), which controls for the effect of the treatment on dispersion. We report the µ value, which is the coefficient used to estimate relative abundance in the *corncob* model and is proportional to the fold-change in relative abundance between the treatment and control. We also assessed differential abundances of taxa in the microaggregate fractions as compared to the bulk soil communities.

### 2.6 Community assembly process assignment

In order to determine the influential community assembly processes within each treatment and fraction, we first applied the method developed by Stegen et al. (2012, 2013, 2015), which compared sample pairs of interest (each possible pair of samples from the same site, tillage treatment, and fraction) to null models that represent stochastic assembly in order to determine the relative influence of selection (based on phylogenetic distances), or dispersal (based on taxonomic dissimilarities). Briefly, the influence of selection was first tested using the abundance-weighted beta-mean nearest taxon distance (βMNTD; the mean phylogenetic distance between each OTU in one community and its closest relative in another community) (*picante::comdistnt*) (Kembel et al., 2010). Homogeneous selection was identified in comparisons for which βMNTD was more than 2 standard deviations below the mean of the null distribution, indicating lower mean phylogenetic distance between pairwise communities than observed in the null. Variable selection was identified in comparisons for which βMNTD was more than 2 standard deviations above the mean of the null distribution, indicating higher mean phylogenetic distance between pairwise communities than observed in the null. Comparisons that fell within 2 standard deviations of the null mean were considered to lack a dominant influence of selection, and were subsequently tested for the influence of dispersal using the modified Raup-Crick metric based on Bray–Curtis dissimilarities (RC_Bray_) (*phyloseq::distance*) (Chase et al., 2011). Homogenizing dispersal was identified in comparisons for which RC_Bray_ was significantly lower than the mean of the null distribution, indicating a higher level of similarity between community compositions than was observed in the null condition; and dispersal limitation was identified in comparisons for which RC_Bray_ was significantly higher than the null mean, indicating lower similarity. Comparisons that were similar to the null mean for both metrics were considered undominated by any particular community assembly process. This analysis, including null model construction, was first performed for each site, and then separately for sets of samples representing each site × treatment × fraction combination.

In addition to determining the dominant community assembly processes based on comparison of the full communities represented by each sample, we also assigned community assembly processes at a finer taxonomic scale, by separately assessing phylogenetically-related bins of OTUs (*iCAMP::pdist.big* and *iCAMP::icamp.big*), as detailed by (Ning et al., 2020). This approach is analogous to the Stegen method, but captures the various ecological mechanisms governing community assembly of the microbial subcommunities within each soil sample. Like the above-detailed full community assessment, here we also compared βMNTD and RC_Bray_ metrics to null models to determine the influences of selection and dispersal, respectively, though sample-to-sample comparisons were made within phylogenetic bins, and the dominant process was weighted by that bin’s relative abundance. We selected bins and ran the analysis using the default parameters as detailed in Ning et al. (2020) and the R documentation (i.e., minimum of 24 OTUs per bin, confirmed by phylogenetic signal testing using *iCAMP::dniche* and *iCAMP::ps.bin*; phylogenetic null model randomization within bins; taxonomic null model randomization across all bins), with the exception of the phylogenetic distance metric, for which we used βMNTD so that results would be more comparable to the full-community scale assessment based on the method by Stegen et al. (2012, 2013, 2015). To test for a significant effect of tillage treatment on the influence of community assembly processes that had > 5% influence, we performed ANOVA as described above. Community relative abundance data was Hellinger-transformed for all community assembly assessments (Legendre and Gallagher, 2001).

### 2.7 Exclusion of plots from analysis

At the Lancaster site, one no-tillage plot and one tillage plot were excluded from analysis. Though this field has been under long-term tillage treatments since the early 1990’s, there are strong indicators in the dataset that the two plots in question were subjected to treatments or conditions that differentiate them from the other plots, likely due to the split-plot use of manure and/or corn fungicide treatments over the years, or the disruptive installation of large pan lysimeters. Though we made every attempt to avoid areas where manure was applied or lysimeters were installed, it was challenging to confirm the precise boundaries of historic split-plots, and the history of fungicide application (unpublished) was unbeknownst to us prior to sampling. The microbial communities of the excluded plots are clearly differentiated in the PCoA (Fig. S1).

## 3. Results

### 3.1 Tillage generally decreased aggregation, with different site responses

At Arlington, over 60% of soil (dry mass basis) was in water-stable aggregate fractions (Fig. 2A), with over 50% of soil in the macroaggregate fraction. There were no significant differences in proportion of soil in macroaggregate, free microaggregate, or silt + clay fractions due to tillage treatment, but within the macroaggregate fraction (Fig. 2C) there was a significant decrease in proportion of soil in the occluded microaggregate fraction (*p* < 0.001; Fig. 2C) from 28% in no-tillage to 22% with tillage, with a complementary increase in the occluded silt + clay fraction (*p* < 0.001).

**Figure 1.**
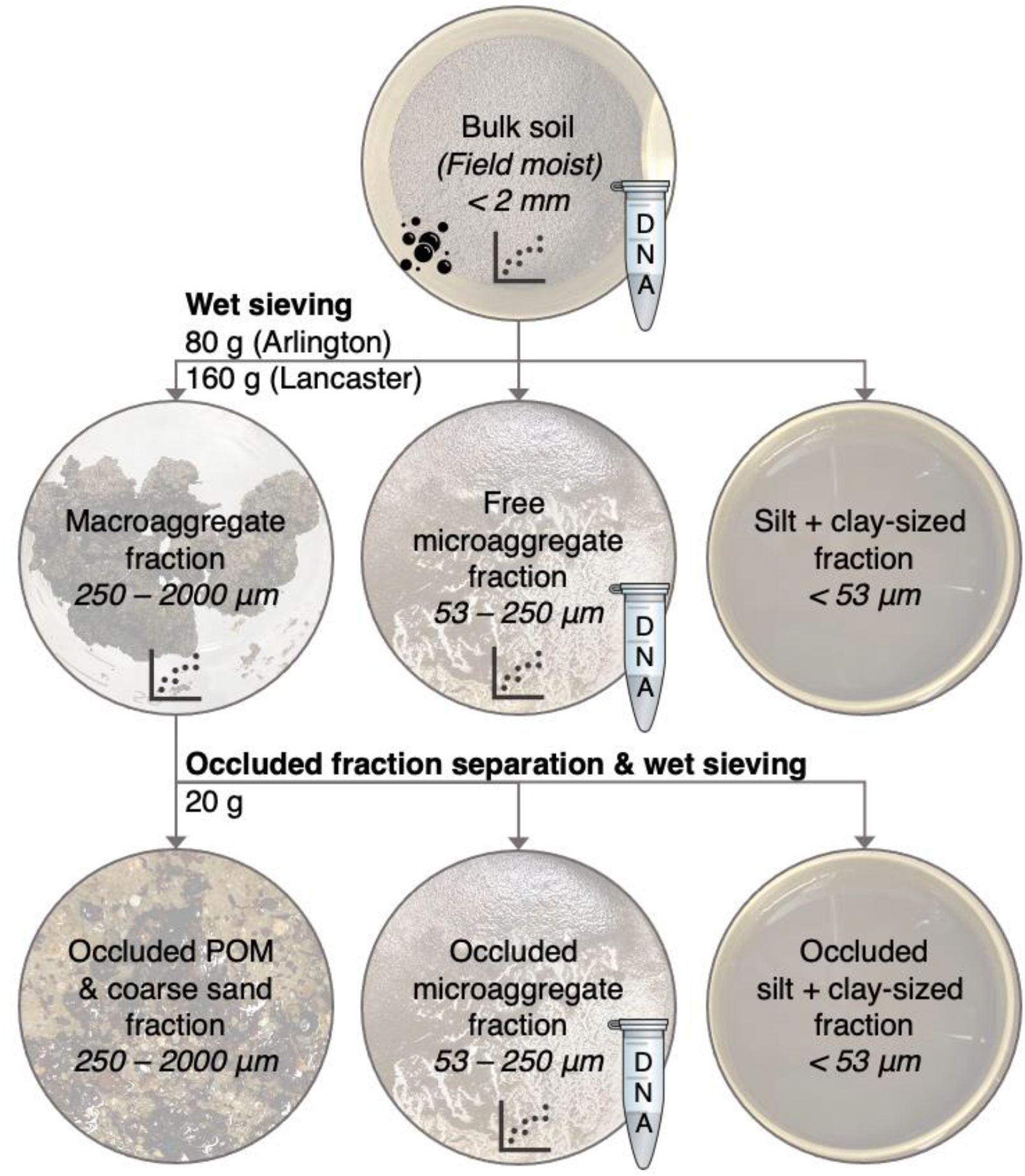
Aggregate size fractionation schematic. Bulk soil (80 g or 160 g) was subjected to wet sieving to separate macroaggregate (250*–*2000 µm), free microaggregate (53–250 µm), and silt + clay-sized (< 53 µm) fractions. A 20 g (wet) subsample of the macroaggregate fraction was then further separated into occluded microaggregate (53–250 µm), occluded silt + clay-sized (< 50 µm), and occluded POM + coarse sand fractions (250–2000 µm). The “DNA” tube indicates that subsamples were retained for 16S rRNA gene amplicon sequencing. The graph icon indicates that subsamples were collected to measure total carbon and total nitrogen. The bubble icon indicates that soil respiration was measured on bulk soil.

**Figure 2.**
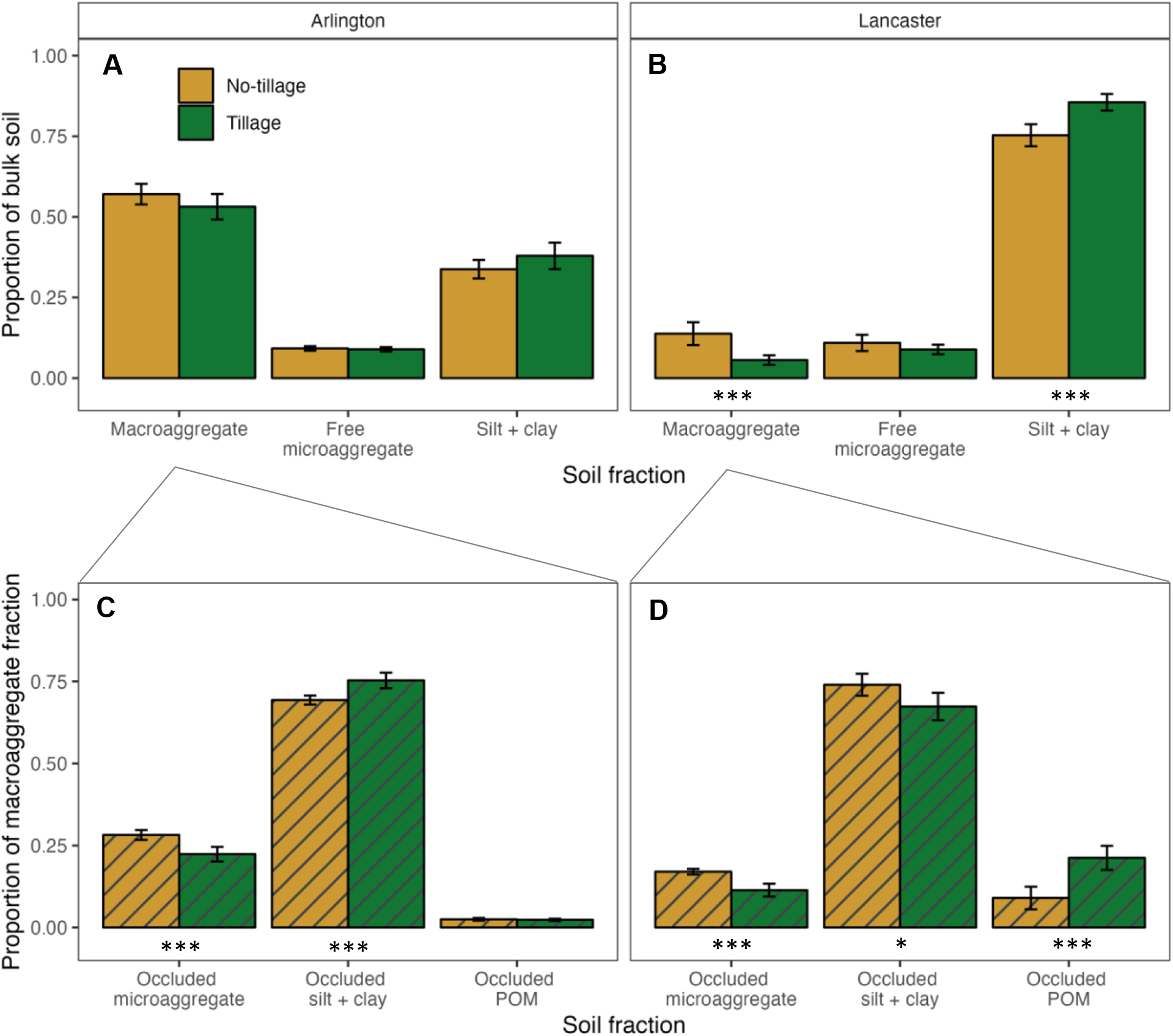
Distribution of bulk soil in various fractions at Arlington, WI (**A**) and Lancaster, WI (**B**), on a dry soil basis. Lower panels show distribution of macroaggregate soil in the occluded fractions (**C** and **D**). Macroaggregate = macroaggregate fraction, 250–2000 µm; Free microaggregate = microaggregate fraction from bulk soil, 53–250 µm; Silt + clay = silt and clay-sized fraction from bulk soil, <53 µm; Occluded microaggregate = microaggregate fraction occluded within macroaggregate fraction, 53–250 µm; Occluded silt + clay = silt and clay-sized fraction occluded within the macroaggregate fraction, <53 µm; Occluded POM = particulate organic matter and sand occluded within the macroaggregate fraction, 250–2000 µm. Error bars represent ± 1.96 SE. Asterisks indicate significant tillage treatment differences within soil fraction: *** = *p* < 0.001, ** = *p* < 0.01, * = *p* < 0.05. Striped bars represent occluded fractions.

At Lancaster, soil was largely unaggregated, with 25% and 14% of soil (dry mass basis) in water-stable aggregate fractions in the no-tillage and tillage treatments, respectively (Fig. 2B). In particular, the tillage treatment had a significantly lower proportion of soil in the macroaggregate fraction, with 6% as compared to 14% in the no-tillage treatment (*p* < 0.001). This was complemented by a significantly higher proportion in the silt + clay fraction (*p* < 0.001). Within the macroaggregate fraction at Lancaster (Fig. 2D), the proportions of the occluded microaggregate fraction and occluded silt + clay fractions were both significantly lower with tillage as compared to no-tillage (*p* < 0.001 and *p* < 0.05, respectively), with 17% of macroaggregate soil in the occluded microaggregate fraction in no-tillage, down to 11% with tillage. There was also a significant increase in occluded POM (*p* < 0.001), from 9% in the no- tillage treatment to 21% with tillage.

### 3.2 Tillage reduced total soil carbon

Tillage decreased total carbon content in all measured fractions at both sites, reported here on a per unit of bulk soil basis (Fig. 3; *p* < 0.001 for each fraction at Arlington, *p* < 0.05 for each fraction at Lancaster).

**Figure 3.**
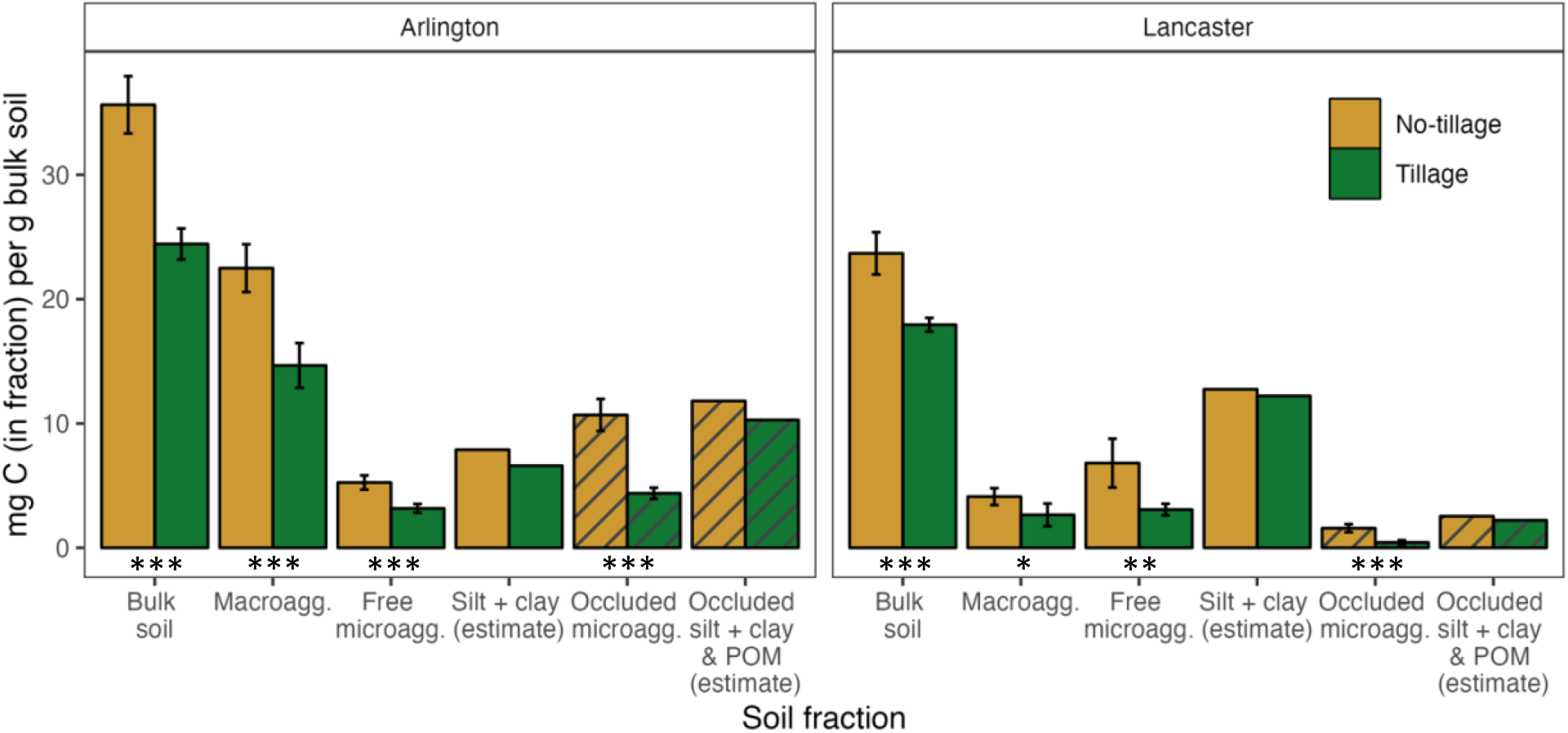
Carbon content of each soil fraction, on a per unit bulk soil basis. Bulk soil = whole soil; Macroagg. = macroaggregate fraction, 250–2000 µm; Free microagg. = microaggregate fraction from bulk soil, 53–250 µm; Silt + clay (estimate) = Carbon content in the < 53 µm fraction, estimated as Bulk soil – (Macroagg. + Free microagg.); Occluded microagg. = microaggregate fraction occluded within macroaggregate fraction, 53–250 µm; Occluded silt + clay & POM (estimate) = Carbon content in the < 53 µm fraction occluded within the macroaggregate fraction, estimated as Macroagg. – Occluded microagg. Error bars represent ± 1.96 SE. Asterisks indicate significant treatment differences within soil fraction: *** = *p* < 0.001, ** = *p* < 0.01, * = *p* < 0.05. The estimated silt + clay carbon contents do not have associated error bars or statistics. Striped bars represent occluded fractions.

Soil C concentration was affected by tillage treatment (Arlington only, *p* < 0.001), soil fraction (both sites, *p* < 0.001), and had a significant interaction effect (both sites, *p* < 0.001) (Table 1 and Fig. S3). At Arlington, C concentration decreased with tillage, overall and within each fraction (*p* < 0.001, Tukey’s HSD). Comparing aggregate fractions, C concentrations of both free and occluded microaggregate fractions were greater than those of the bulk soil and macroaggregate fraction (*p* < 0.05, Tukey’s HSD) in both treatments, and C concentration of the occluded microaggregate fraction was greater than the free microaggregate fraction in no-tillage only (*p* < 0.01, Tukey’s HSD). At Lancaster, C concentration decreased from occluded microaggregate > free microaggregate > macroaggregate > bulk soil (*p* < 0.01, 0.05, and 0.001, respectively, Tukey’s HSD). Comparing aggregate fractions, the C concentrations of both free and occluded microaggregate fractions were greater than C concentrations in the macroaggregate fraction and the bulk soil in the no-tillage treatment only (*p* < 0.001, each comparison, Tukey’s HSD), and C concentration of the free microaggregate fraction was greater in no-tillage as compared to tillage (*p* < 0.001, Tukey’s HSD).

**Table 1.** Soil carbon and nitrogen concentrations and C:N ratio in soil fractions, by tillage treatment, presented as mean (and standard error) with ANOVA p values. Means followed by the same letter (within site) are not statistically different. Bulk soil = whole soil; Macroaggregate = 250–2000 µm; Free microaggregate = microaggregate fraction from bulk soil, 53–250 µm; Occluded microagg. = microaggregate fraction occluded within macroaggregate fraction, 53–250 µm.

|  | *Soil fraction* | Arlington |  | Lancaster |  |
| --- | --- | --- | --- | --- | --- |
|  |  | No-tillage | Tillage | No-tillage | Tillage |
| Soil total carbon concentration (% C) | Bulk soil | 3.56 (0.12) c | 2.44 (0.06) d | 2.37 (0.09) DE | 1.79 (0.03) E |
|  | Macroaggregate | 3.95 (0.14) c | 2.73 (0.09) d | 3.14 (0.16) CDE | 4.90 (0.58) BC |
|  | Free microaggregate | 5.72 (0.23) b | 3.55 (0.15) c | 6.38 (0.61) AB | 3.67 (0.45) CD |
|  | Occluded microagg. | 6.67 (0.32) a | 3.75 (0.09) c | 6.94 (0.32) A | 6.35 (0.55) AB |
| *Source of variation* |  | *p* value |  | *p* value |  |
| Tillage treatment |  | < 0.001 *** |  | 0.0713 |  |
| Soil fraction |  | < 0.001 *** |  | < 0.001 *** |  |
| Tillage × fraction |  | < 0.001 *** |  | < 0.001 *** |  |
| Soil total nitrogen concentration (% N) | Bulk soil | 0.288 (0.007) c | 0.212 (0.004) d | 0.240 (0.010) BC | 0.186 (0.003) C |
|  | Macroaggregate | 0.323 (0.009) c | 0.237 (0.006) d | 0.305 (0.015) B | 0.313 (0.026) B |
|  | Free microaggregate | 0.420 (0.014) b | 0.284 (0.009) c | 0.478 (0.039) A | 0.266 (0.032) BC |
|  | Occluded microagg. | 0.498 (0.018) a | 0.307 (0.006) c | 0.578 (0.022) A | 0.476 (0.036) A |
| *Source of variation* |  | *p* value |  | *p* value |  |
| Tillage treatment |  | < 0.001 *** |  | < 0.001 *** |  |
| Soil fraction |  | < 0.001 *** |  | < 0.001 *** |  |
| Tillage × fraction |  | < 0.001 *** |  | < 0.001 *** |  |
| C:N ratio | Bulk soil | 12.4 (0.15) b | 11.5 (0.12) c | 9.9 (0.32) D | 9.7 (0.22) D |
|  | Macroaggregate | 12.2 (0.16) b | 11.5 (0.13) c | 10.5 (0.68) CD | 15.3 (0.62) A |
|  | Free microaggregate | 13.6 (0.18) a | 12.5 (0.16) b | 13.3 (0.34) B | 13.7 (0.25) AB |
|  | Occluded microagg. | 13.3 (0.19) a | 12.2 (0.13) b | 12.0 (0.21) BC | 13.3 (0.33) B |
| *Source of variation* |  | *p* value |  | *p* value |  |
| Tillage treatment |  | < 0.001 *** |  | < 0.001 *** |  |
| Soil fraction |  | < 0.001 *** |  | < 0.001 *** |  |
| Tillage × fraction |  | 0.428 |  | < 0.001 *** |  |

Soil N concentration trends were very similar to those of total C (Table 1), where N concentration at Arlington decreased with tillage for all soil fractions (*p* < 0.001, all fractions, Tukey’s HSD), and N concentrations of the free and occluded microaggregate fractions were greater than the macroaggregate fraction or bulk soil concentrations (*p* < 0.001, all comparisons, Tukey’s HSD). At Lancaster, N concentrations in the microaggregate fractions were greater than the macroaggregate fraction and bulk soil concentrations for the no-tillage treatment (*p* < 0.001, all comparisons, Tukey’s HSD). The N concentration of the free microaggregate fraction decreased by almost half with tillage (*p* < 0.001, Tukey’s HSD).

The soil C:N ratio demonstrated significant effects of tillage treatment (*p* < 0.001) and soil fraction (*p* < 0.001) at both sites (Table 1 and Fig. S4). At Arlington, C:N ratio was greater in no-tillage compared to tillage (*p* < 0.001, Tukey’s HSD), and greater in free and occluded microaggregate fractions compared to the macroaggregate fraction or bulk soil (*p* < 0.001, Tukey’s HSD). At Lancaster, there was a significant interaction effect of tillage and soil fraction (*p* < 0.001), with a significantly higher C:N ratio in the macroaggregate fraction with tillage.

Tillage decreased respiration (CO_2_ evolution from sieved, field-moist bulk soil) by 50% at Arlington (*p* < 0.01; Fig. S5) on a *per unit soil* basis, was though this difference was not significant on a *per unit soil C* basis (*p* = 0.106). Tillage did not have a significant effect on respiration at Lancaster. No-tillage plot samples averaged 23% soil moisture at both sites, whereas tillage plots averaged 19–20% soil moisture; no adjustments to soil moisture were made prior to respiration measurements.

Soil pH varied significantly by fraction (*p* < 0.001 for both sites; Fig. S6), with bulk soil > occluded microaggregate ≥ free microaggregate fraction, and mean pH of 7.1, 6.8, 6.5 (Arlington) and 7.2, 6.9, and 6.9 (Lancaster), respectively. Lancaster pH data also demonstrated a significant interaction between tillage treatment and soil fraction (*p* < 0.001); bulk soil pH in no-tillage was higher than with tillage (7.4 vs. 7.1), whereas in the occluded microaggregate fraction, soil pH in no-tillage was lower than with tillage (6.5 vs. 7.3), with no significant difference in pH between treatments within the free microaggregate fraction (Fig. S6).

### 3.3 Tillage affected bacterial community composition, but not richness

Bacterial community composition was significantly affected by tillage treatment at both sites (Fig. 4; R^2^ = 0.30 and *p* < 0.001 at Arlington, R^2^ = 0.22 and *p* < 0.001 at Lancaster; PERMANOVA). The homogeneity of variance test (BETADISPER) was also significant for tillage treatment at Arlington and Lancaster (*p* < 0.001, *p* < 0.05, respectively), which indicates that the assumptions of the PERMANOVA were not met due to differences in sample dispersion.

**Figure 4.**
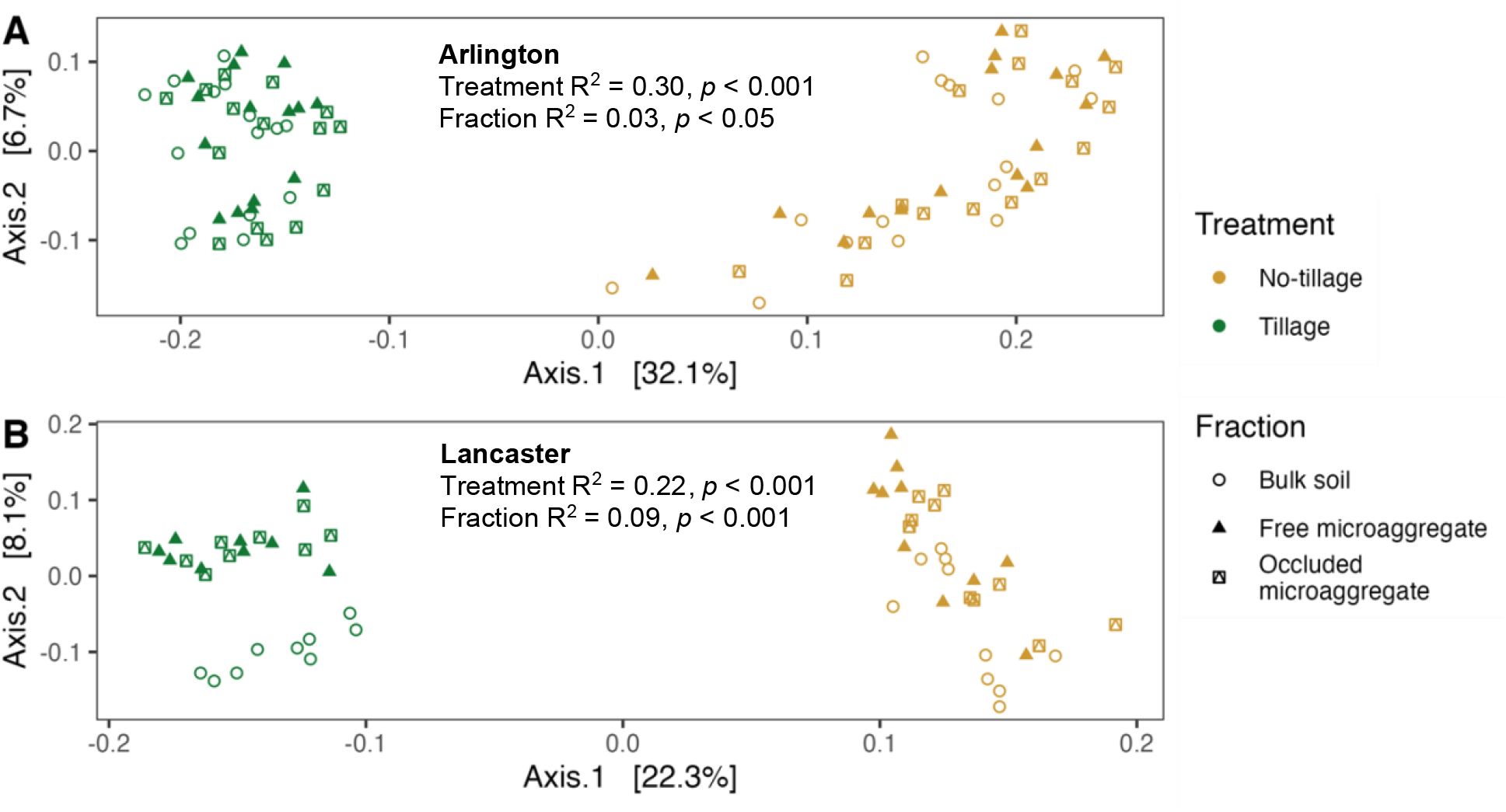
Principal coordinates analysis of Bray-Curtis dissimilarities of Hellinger-transformed community relative abundances, by tillage treatment for Arlington, WI (**A**), and Lancaster, WI (**B**). Each point represents the community of one sample-fraction. Soil fractions are as follows: Bulk soil = whole soil; Free microaggregate = microaggregate fraction from bulk soil, 53–250 µm; Occluded microaggregate = microaggregate fraction occluded within macroaggregate fraction, 53–250 µm. Displayed statistics are from PERMANOVA.

Tillage decreased dispersion of community composition by 14% and 6% relative to the no-tillage treatment at Arlington and Lancaster, respectively, as quantified by between-plot mean distance to spatial median (Fig. 5A and D; *p* < 0.001 for Arlington and *p* < 0.01 for Lancaster). This trend, which indicates higher dissimilarity of samples within the no-tillage treatment, was also apparent at the plot scale, where tillage decreased sample dispersion within plots by 13% and 5% relative to no-tillage at Arlington and Lancaster, respectively (Fig. 5B and E; *p* < 0.001 for Arlington and *p* < 0.05 for Lancaster). The dispersion of the free microaggregate and occluded microaggregate communities within each soil core did not significantly differ between tillage treatments, though there was a trend towards decreased dispersion with tillage at Lancaster (*p* < 0.1; Fig. 5C and F). There were no significant differences in richness estimates (Fig. S7) or Faith’s PD (Fig. S8) attributable to tillage treatments at either site.

**Figure 5.**
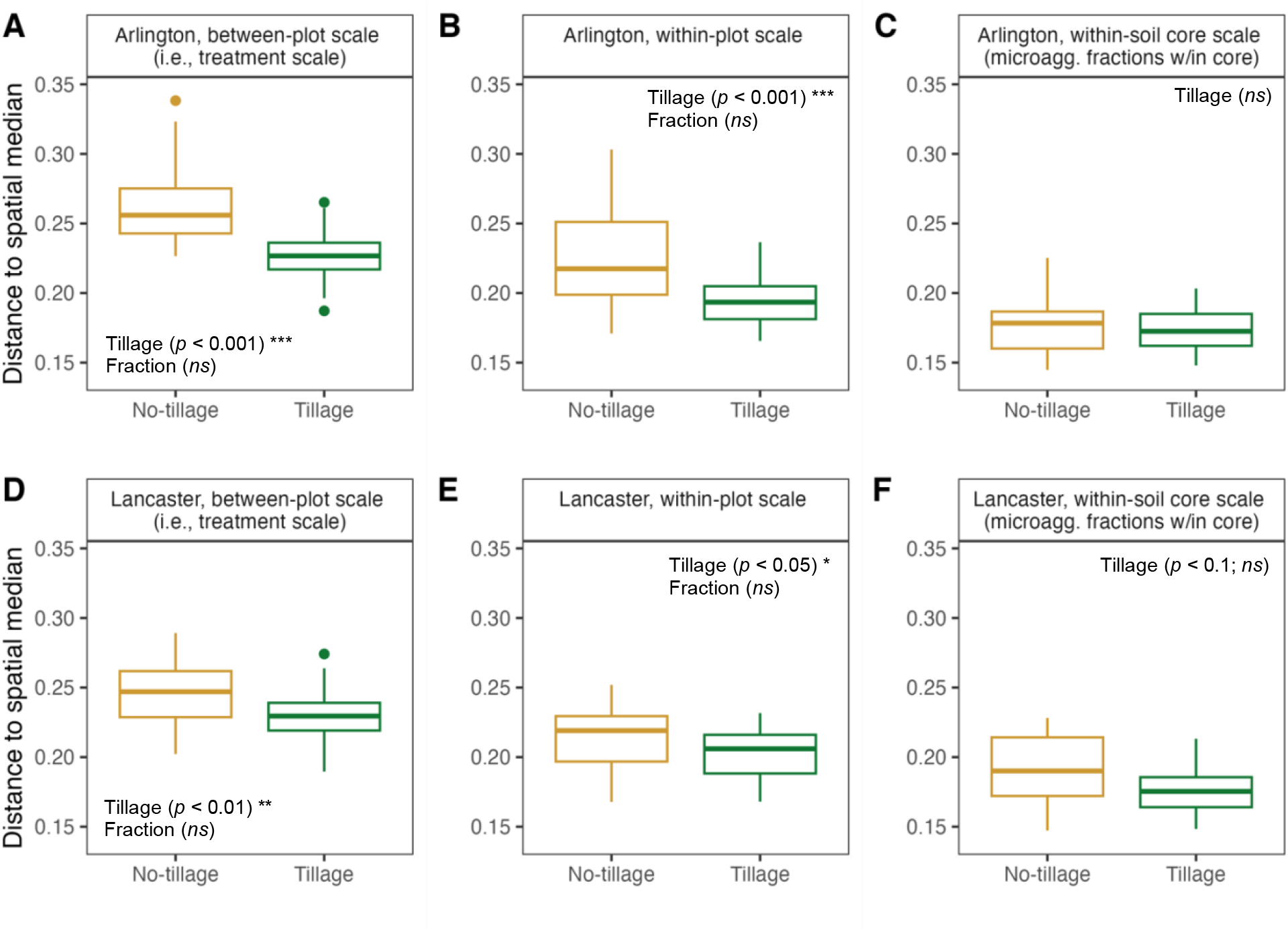
Dispersion of community compositional data as Bray-Curtis dissimilarities, represented as distance to spatial median at the between-plot scale (i.e., treatment scale; **A** and **D**); within-plot scale (**B** and **E**); and soil core scale (free vs. occluded microaggregate fraction samples within a soil core; **C** and **F**) at Arlington, WI (**A**, **B**, and **C**) and Lancaster, WI (**D**, **E**, and **F**). Data presented in A, B, D, and E represent bulk soil and both microaggregate fractions together.

### 3.4 Community composition differed slightly between microaggregate fractions

There was a significant effect of soil fraction on bacterial community composition at both sites (R^2^ = 0.03 and *p* < 0.05 for Arlington and R^2^ = 0.09 and *p* < 0.001 for Lancaster, PERMANOVA; Fig. 4). Pairwise testing demonstrated significant differences between bulk soil and the free microaggregate fraction, and between bulk soil and the occluded microaggregate fraction at Lancaster only (*p* < 0.01), whereas pairwise testing amongst soil fractions was not significant at Arlington. Sample dispersion was homogeneous (i.e., beta diversity was similar) across soil fractions at both treatment and plot scales at both sites (Fig. 5). There was no interaction effect of tillage treatment × soil fraction on community composition at either site.

Richness estimates demonstrated a significant effect of soil fraction (*p* < 0.05, Fig. S7) at Lancaster only; the richness estimate for the occluded microaggregate fraction was 8% lower than that of the bulk soil (*p* < 0.05, Tukey’s HSD). Faith’s PD was also affected by fraction (*p* < 0.05, Arlington, and *p* < 0.001, Lancaster; Fig. S8) by which the occluded microaggregate fraction was significantly lower than that of bulk soil at both sites (*p* < 0.05 and *p* < 0.001, respectively, Tukey’s HSD), and the free microaggregate fraction was also lower than bulk soil at Lancaster (*p* < 0.01, Tukey’s HSD).

### 3.5 Weighted mean predicted 16S rRNA gene copy number increased with tillage

At Arlington, there was small but statistically significant 7% increase in the weighted mean predicted 16S rRNA gene copy number with tillage (*p* < 0.001; Fig. 6). Fraction was also significant (*p* < 0.001), and there was a significant interaction effect of tillage and fraction (*p* < 0.05). The weighted mean predicted 16S gene copy number was lower in the occluded microaggregate fraction relative to the bulk soil or free microaggregate fraction in the tillage treatment, whereas weighted mean predicted 16S gene copy number was similar across fractions of the no-tillage treatment. At Lancaster, there was a significant 10% increase in the weighted mean predicted 16S gene copy number with tillage (*p* < 0.001), and no significant effect of fraction or interaction effect.

**Figure 6.**
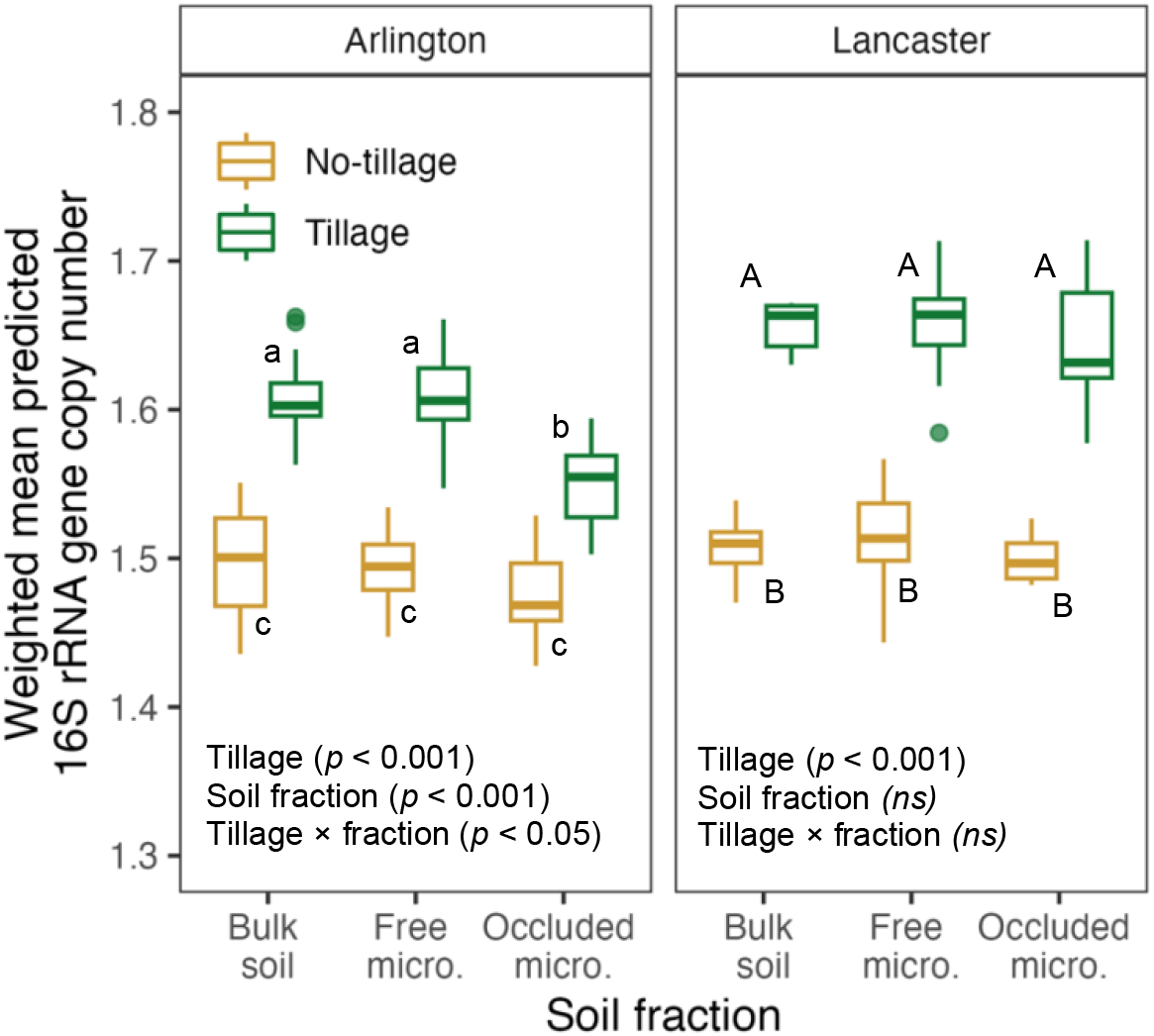
Weighted mean predicted 16S rRNA gene copy number. These data represent taxa for which a gene copy number was available in the rrnDB (Stoddard et al., 2015). Bulk soil = whole soil; Free micro. = microaggregate fraction from bulk soil, 53–250 µm; Occluded micro. = microaggregate fraction occluded within macroaggregate fraction, 53–250 µm. Boxplots with the same letter (within site) are not statistically different.

### 3.6 Influence of homogenizing dispersal increases with tillage

Using the full community comparison approach, community assembly was dominated by homogeneous selection, with > 95% of pairwise comparisons across each set of samples (representing site, tillage treatment, and fraction combinations), when null models were comprised of site-wide data. This outcome is consistent with the intensity and uniformity of management in monocrop systems (Figuerola et al., 2015).

We next performed the community-wide approach separately for each set of samples (site × treatment × fraction combination) in order to further scrutinize community assembly processes. This approach, critically, employed narrower null models that were more specific to the pairwise comparisons. The results from that analysis demonstrate evidence for homogenizing dispersal amongst other processes, while still emphasizing the importance of homogeneous selection in these systems (see Fig. S9, for full results using treatment-specific null models).

Going forward, and in the Discussion section, we will focus on the results from the OTU binning-based approach (Ning et al., 2020), which indicated that homogeneous selection had a ∼14% relative influence across treatments and fractions, for both within-plot and between-plot comparisons at Arlington (Fig. 7A and B), with the between-plot comparisons demonstrating a significant decrease in homogeneous selection under tillage relative to no-tillage (*p* < 0.05). The influence of homogenizing dispersal significantly increased with tillage (*p* < 0.001), from 25% to 46% in bulk soil for within-plot comparison; and from 12% to 28% for between-plot comparisons. There was also a large proportion of undominated comparisons— 30–60% at the within-plot scale and 50–70% at the between-plot scale.

**Figure 7.**
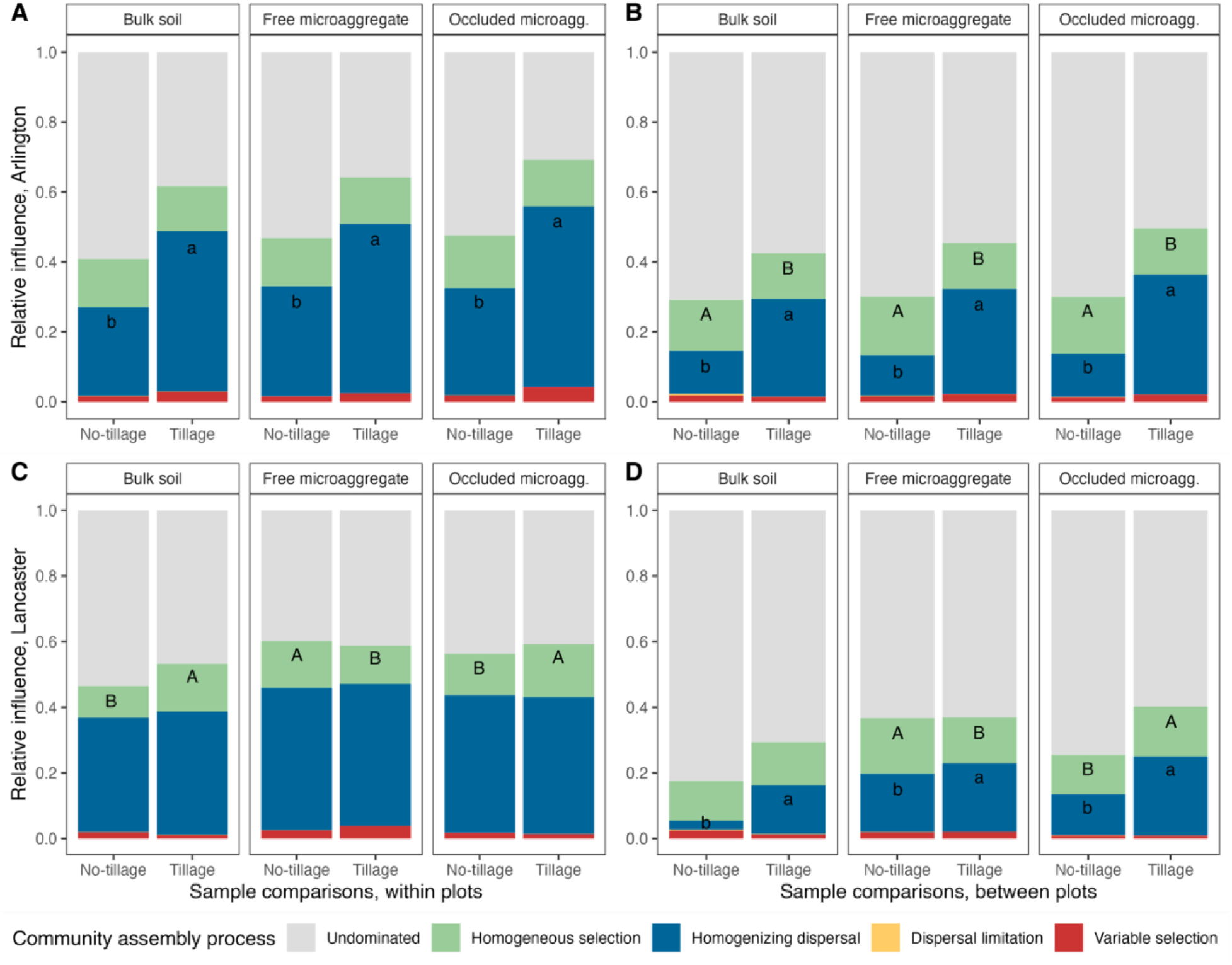
The dominant community assembly processes, by tillage treatment, within bulk soil, free microaggregate, and occluded microaggregate fractions at Arlington, WI (**A** and **B**); and Lancaster, WI (**C** and **D**). Sample comparisons were made within-plot (**A** and **C**) or between-plot (**B** and **D**). Community assembly processes were assigned within phylogenetically related bins of OTUs for pairwise comparisons of samples using a null modeling approach, and weighted by the relative abundance of OTUs in that bin to comprise the full community (Ning et al., 2020). As detailed in the text, first the influence of selection was determined using the *β*-mean nearest taxon distance, and then the influence of dispersal was determined using the modified Raup- Crick metric based on Bray-Curtis dissimilarity. For homogeneous selection and homogenizing dispersal (the processes with > 5% influence), different letters signify a statistically significant difference in the influence of that process due to tillage (within site and fraction).

When using the OTU binning-based approach at Lancaster (Fig. 7C and D), the within-plot comparisons demonstrated similar trends as those at Arlington regarding influences of homogeneous selection (∼15%), homogenizing dispersal (∼40%), and undominated (∼45%). Unlike Arlington, there was no significant effect of tillage on the influence of homogenizing dispersal at the within-plot scale, whereas homogenizing selection significantly increased from 10% to 15% of relative abundance in the bulk soil (*p* < 0.05). In the between-plot comparisons, the influence of homogenizing dispersal significantly increased with tillage in the bulk soil and occluded microaggregate fraction (*p* < 0.001), whereas homogeneous selection experienced a small decrease with tillage in the free microaggregate fraction, and a small increase with tillage in the occluded microaggregate fraction (*p* < 0.05, and *p* < 0.01, respectively). For the between- plot comparisons, most were undominated— 60–80%.

### 3.7 Taxonomic differences attributable to tillage

The most common phyla across the datasets were *Actinobacteria*, *Acidobacteria*, and *Proteobacteria*, which together comprised about 60% of all reads at each site in both tillage treatments. At Arlington, there was generally a consistent phylum-level response across the bulk soil, free microaggregate, and occluded microaggregate fraction by which tillage resulted in significant increases in relative abundances of *Actinobacteria*, *Armatimonadota*, *Chloroflexi*, *Cyanobacteria*, *Firmicutes*, *Gemmatimonadetes*, and *Methylomirabilota*, and significant decreased relative abundances of *Acidobacteria*, *Myxococcota*, *Proteobacteria*, and *Verrucomicrobia* (see Fig. S10 for relative abundances and *p* values). At Lancaster, tillage resulted in significant increases in relative abundances of *Chloroflexi*, *Cyanobacteria*, and *Gemmatimonadetes*, and significant decreases in relative abundances of *Crenarchaeota* and *Verrucomicrobia* (see Fig. S11 for relative abundances and *p* values).

We also identified key taxa associated with no-tillage and tillage treatments, based on differential abundance. Across both sites, we identified a total of 1658 taxa that were enriched with tillage (relative to no-tillage), and 1602 taxa that were enriched in no-tillage (relative to the tillage treatment). See Supplementary Table S2 for a complete list of enriched taxa, with coefficients of differential abundance (µ) and sequences. We focused on the taxa with the biggest responses (µ > 1.0), and only considered enriched taxa with mean relative abundances greater than 0.001 (0.1%), which resulted in 15 and 9 focal taxa enriched with tillage and no-tillage, respectively, at Arlington, and 9 and 4 focal taxa enriched with tillage and no-tillage, respectively, at Lancaster (Figs. S12 and S13). Though some taxa were unique responders within a soil fraction, there were numerous taxa that were enriched across bulk soil, free microaggregate, and occluded microaggregate fractions.

### 3.8 Taxonomic differences between microaggregate fractions

We also identified taxa that were enriched in the microaggregate fractions using differential abundance. We identified a total of 382 taxa across both sites that were enriched in the free or occluded microaggregate fractions relative to the bulk soil. See Supplementary Table S3 for a complete list of enriched taxa, with coefficients of differential abundance (µ) and sequences. Narrowing our focus on taxa with the biggest responses (µ > 1.0), and not overly rare (mean relative abundances greater than 0.001 [0.1%]), there were 8 and 10 taxa enriched in the free microaggregate and occluded microaggregate fractions at Lancaster, respectively, most of which were in the tillage treatment (Fig. S14), and no taxa that fit those parameters at Arlington. There were several *Chloroflexi* OTUs representing the class *Anaerolineae*, and several *Cyanobacteria* OTUs that were relatively enriched in the occluded microaggregate fraction. We did not assess taxa that were depleted in microaggregate communities relative to the bulk soil because the former is inherently a subset of the latter.

## 4. Discussion

We examined the effects of tillage on soil bacterial community composition and assembly, specifically in the free and occluded microaggregate fractions, and will discuss these findings with respect to soil carbon protection as modulated through changes to soil aggregation. Findings generally supported our hypotheses that tillage would homogenize bacterial communities, with community assembly driven by homogenizing dispersal, but only weakly supported the hypothesized distinctions between the free and occluded microaggregate communities. Overall, we found decreased aggregation, soil C, and soil N with tillage (Figs. 2 and 3, Table 1), which agrees with previous work (Frey et al., 1999; Six et al., 1999; Al-Kaisi et al., 2014; Zheng et al., 2018). This supports the paradigm that tillage increases macroaggregate turnover, thus derailing occluded microaggregate formation, and decreasing soil C content through enhanced decomposition and weakened long-term protection (Six et al., 1999; King et al., 2019).

### 4.1 Tillage decreased soil aggregation and soil carbon

Our work provides further evidence supporting the relationship between soil aggregation and SOC content, while reiterating that tillage reduces aggregation and SOC in surface soil. We found that 90% of the increase in SOC under no-tillage relative to tillage was in aggregate fractions, with the majority (> 75%) of this increase specifically in the microaggregate fractions (Figs. 3 and S2). At Arlington, most of the increase in C under no-tillage was attributed to the occluded microaggregate fraction, which reflects previous work (Denef et al., 2004; Six and Paustian, 2014). However, at Lancaster, most of the increase in C was in the free microaggregate fraction, which could reflect post-season sample timing with respect to macroaggregate seasonal dynamics. As roots and hyphae die following crop plant senescence, macroaggregates rapidly destabilize, liberating previously occluded microaggregates into the free microaggregate pool (Perfect et al., 1990; Oades and Waters, 1991), as detailed below in Section 4.6.

As with the difference in SOC accumulation between the microaggregate fractions, the two sites continue to tell somewhat different stories of aggregation and SOC distribution. Arlington exemplifies the “cultivation loop” (sensu Six et al., 1999), by which tillage stimulates decomposition and macroaggregate turnover, thus precluding SOM enrichment and resulting in older, C-depleted microaggregate fractions (Table 1). Simultaneously, under no-tillage, undisturbed macroaggregates foster development of new occluded microaggregates, as indicated by higher C concentrations and wider C:N ratios in the microaggregate fractions (Table 1).

On the contrary, at Lancaster, the macroaggregate fraction *under tillage* had a high C concentration, wide C:N ratio, and increased proportion of macroaggregate-occluded POM relative to no-tillage (Table 1, Figs. 2 and S4), indicative of largely undecomposed residue. Substantial residue at Lancaster is a testament to the continuous corn rotation; the residue from the previous crop (corn) was potentially double that of Arlington (where the previous crop was soybean), and of a higher C:N ratio (Ordóñez et al., 2020). Tillage breaks up crop residue, bringing it into direct contact with mineral particles and soil microbiota to nucleate new macroaggregates, which could have enhanced C and POM concentration in the macroaggregate fraction, despite the overall tillage-driven decrease in proportion of macroaggregates. Though counter to how we typically characterize macroaggregates under tillage (e.g., low soil C and POM concentrations), this evidence for less processed SOM in the macroaggregate fraction supports the overall narrative of a shorter mean macroaggregate lifespan under higher turnover with tillage (Elliott, 1986). In contrast, the corn-soy rotation at Arlington resulted in more straightforward soil C trends (e.g., C concentration in no-tillage > tillage; C concentration in microaggregates > macroaggregates and bulk soil; Table 1 and Fig S3). It would be relevant to repeat these measurements shortly after a fall tillage event to assess if tillage accelerates the decomposition of occluded POM and decreases SOC in the macroaggregate fraction, particularly in a system such as Lancaster where these metrics were high just prior to a fall tillage event.

The overall weak aggregation at Lancaster (Fig. 2), with less than 15% of soil in aggregates, lends support to a recently proposed paradigm shift that suggests soils under tillage may not be a relevant application of the physicogenic aggregate, but instead represent engineered, loosely arranged soil fragments that largely lack natural biopore networks (Or et al., 2021).

### 4.2 Tillage homogenized bacterial communities via dispersal

Tillage had a significant effect on bacterial community composition at both sites (Fig. 4), as observed by others (Srour et al., 2020; Bhattacharyya et al., 2021), which resulted in more homogeneous communities at both within-plot and between-plot scales, confirming hypothesis H3 (Figs. 4 and 5). At the within-plot scale, decreased compositional differences with tillage (Fig. 5) may be driven by homogenizing dispersal at Arlington (Fig. 7A), partially confirming hypothesis H1. At Lancaster, the relatively smaller effect of tillage on community composition (Fig. 4) and community dispersion (Fig. 5) may be attributable to the lack of increased influence of homogenizing dispersal, and only small increases in homogeneous selection (Fig. 7).

At the between-plot scale, we might have expected to see increased influences of homogeneous selection and perhaps dispersal limitation with tillage, because management of these plots is similar, yet they are spatially separated. However, like findings at the within-plot scale, tillage also increased the influence of homogenizing dispersal at the between-plot scale at both sites. Therefore, another tillage-driven mechanism increased the compositional similarity amongst these spatially distinct plots, barring direct organismal dispersal, without increasing phylogenetic similarity (which would have increased homogeneous selection). Because tillage systematically preserves more stable, potentially older microaggregates, we may be observing founder effects that manifest as taxonomic similarity between plots in a field (Rillig et al., 2017). Despite some significant shifts in selection and dispersal, community assembly is largely undominated at the between-plot scale, demonstrating a high level of stochasticity and potential for ecological drift.

Despite homogenizing community composition, tillage did not have a significant effect on bacterial richness (Fig. S7). Previous work has found tillage to have contrasting effects on richness with both neutral and negative findings (Constancias et al., 2013; Smith et al., 2016). The tillage practices used at these sites (fall chisel plow plus spring cultivation) are perhaps too infrequent or mild to affect richness estimates, as previous work has found that richness significantly decreased only in soil disturbed at least biweekly (West and Whitman, 2022). It is also possible that sequencing efforts poorly represented the relative richness of these systems and soil fractions (Bach et al., 2018), though the *betta* model that we used for richness estimation is specifically designed to account for unobserved taxa (Willis et al., 2017).

We did not observe strong influences of dispersal limitation or variable selection under no-tillage, as was hypothesized (H1). This may be attributed to the generally homogeneous soil environment that is characteristic of intensively managed monocrop systems, regardless of tillage practices.

### 4.3 Tillage favors potential for fast growth

Increased weighted mean predicted 16S gene copy number under tillage (Fig. 6) was also noted in a recent global metanalysis (Wilhelm et al., 2023), and is consistent with the idea that pulses of resources (e.g., C liberation or litter incorporation via tillage) select for competitors with fast growth potential in soil bacteria (Schmidt et al., 2018). These studies also found that larger mean estimated genome size correlated with lower soil health ratings and tillage, indicating a need for higher metabolic and regulatory capabilities under environmental instability (Schmidt et al., 2018; Wilhelm et al., 2023). However, the fairly uniform effect on weighted mean predicted 16S gene copy number across soil fractions (Fig. 6), which do differ in chemical composition (Table 1), indicate that physical disturbance may also influence fitness as it relates to other aspects of life history strategy, such as chemical signaling, community goods, or secondary metabolites.

For example, this could point to a scenario by which oligotrophic organisms, which invest heavily in extracellular enzymes, are at a disadvantage when proximity to these metabolites is disrupted under physical disturbance (Junkins et al., 2022), whereas copiotrophic generalists, which are less reliant on these proximity-based life strategies, might be less hindered by physical disturbances.

We might have expected tillage-driven increases in weighted mean predicted 16S gene copy number (Fig. 6) to be accompanied by increases in soil respiration (on a per gram C basis), due to lower carbon use efficiency associated with increased 16S gene copy number (Roller et al., 2016). However, the similar rate of C respiration across tillage treatments at both sites (Fig. S5B) implies that the no-tillage and tillage communities processed C similarly, and/or the small increase in weighted mean predicted 16S gene copy number, although statistically significant, was not biologically relevant for C mineralization. Soil respiration—on a per gram soil basis— did decrease under tillage at Arlington (Fig. S5A), as a function of decreased bulk soil C concentration (Table 1). The no-tillage samples averaged slightly higher soil moisture content, which also may help explain the slight increases in respiration. Though our measurements of CO_2_ evolution from sieved soil do not accurately represent respiration of an intact soil and pore network (Vogel et al., 2022), this analysis still indicates that the C mineralization potential of these soil communities may not be limited by tillage-driven compositional changes.

### 4.4 Taxonomic differences due to tillage

Some broad, phylum-level compositional differences follow archetypical expectations under tillage: *Firmicutes*, generally thought to include many fast-growing copiotrophs, increased in relative abundance with tillage, as was previously observed (Schmidt et al., 2018), whereas *Verrucomicrobia* include numerous oligotrophic taxa (Bergmann et al., 2011), and decreased under tillage (Figs. S10 and S11). *Firmicutes* had higher mean relative abundances in the bulk soil as compared to microaggregate fractions. There were several taxa that responded to tillage representing the genus *Nocardioides* (*Actinobacteria*). A responder to frequent soil disturbance (West and Whitman, 2022), *Nocardioides* has been negatively correlated with soil health in a global metanalysis (Wilhelm et al., 2023). We also found relative enrichment of *Sphingomonas* and *Geodermatophilus* under tillage, both of which have been identified as key tillage responders (Wilhelm et al., 2023). Under no-tillage, we also found enrichment of the genus *Gaiella*, (*Actinobacteria),* which was one of several identified bioindicators of high biological soil health ratings (Wilhelm et al., 2023). We also found enrichment of anaerobic taxa (e.g., *Anaerolineae*) (Yamada and Sekiguchi, 2020) in microaggregates (Fig. S14), which have anoxic microsites (Sexstone et al., 1985).

The enrichment of *Cyanobacteria* with tillage (Fig. S12) reflects how repeated disturbance to the soil surface suspends biocrust communities in an early successional stage (Belnap and Eldridge, 2001). Because *Cyanobacteria* are particularly enriched in microaggregate fractions, where we would not expect photosynthetic organisms to survive or thrive, this enrichment may reflect the presence of relic DNA (Carini et al., 2016) or dormant organisms (Lennon and Jones, 2011), integrated from biocrusts into the soil matrix via tillage, and subsequently protected within microaggregates. The specific *Cyanobacteria* taxa enriched in our study (*Microcoleus* PCC-7113 and *Tychonema* CCAP 1459-11BA) were both previously found in soil under organic management, perhaps also due to frequent disturbance (Santoni et al., 2022).

### 4.5 Evidence for fluidity between the free and occluded microaggregate fractions

Within tillage and no-tillage treatments, the bacterial compositions and community assembly patterns of the free and occluded microaggregate fractions are fairy indistinct from each other (*p* > 0.05, PERMANOVA; Figs. 4 and 7), contrary to hypothesis H2. This indicates that these operationally defined fractions likely have substantial overlap, potentially attributable to wholesale shifts of occluded microaggregates to the free fraction at the end of the temperate annual cropping season, when macroaggregates rapidly degrade with root senescence (Oades and Waters, 1991), as detailed in Section 4.6. Further, the community compositional dispersion of free and occluded microaggregate fractions within each soil core was unaffected by tillage treatment (Fig. 5C and F), which indicates that the fluidity between the free and occluded microaggregate fractions may not be particularly responsive to disturbance. Since we did not identify bacterial drivers that explain enhanced SOC persistence in the occluded microaggregate fraction, future work could instead focus on the physical and chemical drivers of C storage and persistence in microaggregate fractions (Bailey et al., 2019; Kravchenko et al., 2019), or fungal community drivers (Lehmann and Rillig, 2015).

Overall, microaggregate communities were less phylogenetically diverse than bulk soil at both sites (Fig. S8), implying that the microaggregate microenvironment selects for a community adapted to that environment. This is consistent with the PERMANOVA results that demonstrated significant compositional differences between the bulk and microaggregate fractions, though these differences were very small (R^2^ = 0.09 at Lancaster and R^2^ = 0.03 at Arlington).

### 4.6 Factors that may have moderated the measured impact of tillage

We will briefly consider several nuanced factors in this study. The tillage treatment at both sites included a fall chisel plowing, which is sometimes considered a reduced or even conservation tillage approach because it is shallower and more moderate compared to moldboard or disk plowing, and does not invert the soil (e.g., Zuber and Villamil, 2016). Previous work has found chisel plow tillage to have the same effect as no-tillage on aggregate stability and microbial biomass (Al-Kaisi et al., 2014; Zuber and Villamil, 2016). Several other factors may obscure or diminish the relative impacts of tillage in this study, including crop-related seasonal macroaggregate dynamics, wet-dry or freeze-thaw cycles, and clay mineralogy.

As noted above, macroaggregates rapidly destabilize following crop senescence—which begins four to eight weeks prior to grain harvest—thus potentially diminishing tillage-driven differences in soil aggregation measured post-harvest (Fig. 2) and liberating occluded microaggregates into the free microaggregate pool (Perfect et al., 1990; Oades and Waters, 1991). Similar aggregation patterns across tillage treatments were previously observed by Huang et al. (2010), in which sampling occurred months after corn harvest. Tillage differences may be further diminished by the physically disruptive effects of freeze-thaw and wet-dry cycles at the soil surface, which would impact aggregate stability of otherwise undisturbed soil under no-tillage (LeGuillou et al., 2012; Bailey et al., 2019). These effects are likely variable in tillage vs. no-tillage treatments, given differences in protective surface residues (Cruse et al., 2001).

Another contributing factor to differences in aggregation and C concentration between sites may be driven by different mineralogy (Denef et al., 2004). Mollisols, such as the soil at Arlington, are generally recognized to promote organo-mineral complexes. The clay mineralogy of the Plano silt loam at Arlington is interstratified smectite-illite (Liu et al., 1997); the high specific surface area of illite may promote SOC retention, and the expansible nature of smectite may physically protect organic matter (Sarkar et al., 2018). The Fayette silt loam at Lancaster (Alfisol) has been mineralogically characterized as predominantly montmorillonite clay minerals, which is also an expansible layer phyllosilicate (Caldwell et al., 1955), though the low activity (1:1) clay may be influential, as evidenced by the very low level of aggregation even under no-tillage (Fig. 2), as was noted for a mixed-mineralogy clay in Six et al. (2000). We find support for these literature-based suppositions in the lower concentrations of base cations and lower overall cation exchange capacity measured at Lancaster (Table S1). These mineralogical differences could explain why Arlington has higher proportions of aggregated soil and higher SOC and SOM concentrations than Lancaster, despite similar texture (silt loam) and relatively similar corn-based cropping systems. Yet another consideration might be the lower sand content across the tillage treatment at Lancaster, with 8% sand in contrast to 18% in the no-tillage treatment, potentially contributing to the relatively high C concentrations in the macroaggregate and occluded microaggregate fractions under tillage, though these sand measurements are somewhat anecdotal because they represent composite samples (n = 1 per treatment). Sand content correction in aggregate fractions could have altered our fraction proportions and soil C concentrations, but this adjustment was not performed because our primary objective for aggregate fractionation was to isolate the microaggregate fractions for bacterial community assessment.

## 5. Conclusions

For the first time, this study demonstrates that tillage homogenizes the soil microbial community through homogenizing dispersal, while supporting previous conclusions that tillage disrupts aggregation and decreases carbon at the soil surface. Counter to our hypothesis, the bacterial communities of the free and occluded microaggregate fractions are highly similar, indicating that microaggregates may readily shift between these operationally defined soil fractions, rather than adhering to purported categories. Tillage may accentuate seasonal changes that characterize temperate annual cropping systems (e.g., crop senescence, freeze-thaw, and wet-dry cycles), which together challenge the strength and longevity of macroaggregates in which occluded microaggregates form and soil carbon is protected. Thus, while our findings reiterate the importance of the occluded microaggregate fraction for soil C persistence, we also suggest that this occluded microaggregate C is subject to an increased rate of turnover upon convergence with the free microaggregate fraction when macroaggregate stability degrades. Conceptually, this underscores how aggregate microhabitats develop and devolve throughout the soil matrix, in concert with microbial activity, forming isolated hotspots driven by resource availability in the patchy soil environment.

## Supplementary Information

Supplementary Information can be found online.

## Acknowledgements

The authors are indebted to the researchers and operators who established and/or maintained these long-term tillage studies over the years, and provided information about their histories, including Thierno Diallo, Doug Wiedenbeck, Satish Gupta, Holly Dolliver, and the crews at the Arlington and Lancaster Agricultural Research Stations. The authors would like to thank Alexa Hanson, Kallysa Taylor, Emma Johnson, and Isabelle Bartholomew for their direct contributions to this project in the lab and field; Erika Marín-Spiotta and members of the Whitman lab for their thoughtful input; Daliang Ning for guidance with iCAMP analysis; and Harry Read and Anna Cates for their perspectives on soil fractionation and use of the microaggregate isolator.

The authors also acknowledge the UW Biotechnology Center DNA Sequencing Facility (Research Resource Identifier—RRID:SCR_017759). Part of this research was performed using the computational resources and assistance of the UW–Madison Center for High Throughput Computing (CHTC) in the Department of Computer Sciences, with the help of Christina Koch. The CHTC is supported by UW–Madison, the Advanced Computing Initiative, the Wisconsin Alumni Research Foundation, the Wisconsin Institutes for Discovery, and the National Science Foundation, and is an active member of the OSG Consortium, which is supported by the National Science Foundation (NSF) and the U.S. Department of Energy’s Office of Science. This work was financially supported by the O.N. Allen Professorship (UW–Madison CALS), the Louis and Elsa Thomsen Wisconsin Distinguished Graduate Fellowship (UW–Madison CALS), and a NSF EAGER grant (award #2024230).

## Author contributions

JW and TW conceived of the project. JL has maintained the tillage experiment in Arlington, WI since 1994. JW collected soil samples, conducted lab work, analyzed the data, and drafted the manuscript. All authors reviewed and edited the manuscript.

## References

Al-Kaisi, M.M., Douelle, A., Kwaw-Mensah, D., 2014. Soil microaggregate and macroaggregate decay over time and soil carbon change as influenced by different tillage systems. Journal of Soil and Water Conservation 69, 574–580. doi:10.2489/jswc.69.6.574

Anderson, M.J., 2006. Distance-Based Tests for Homogeneity of Multivariate Dispersions. Biometrics 62, 245–253. doi:10.1111/j.1541-0420.2005.00440.x

Anderson, M.J., 2001. A new method for non-parametric multivariate analysis of variance. Austral Ecology 26, 32–46. doi:10.1111/j.1442-9993.2001.01070.pp.x

Angers, D.A., Recous, S., Aita, C., 1997. Fate of carbon and nitrogen in water-stable aggregates during decomposition of 13C15N-labelled wheat straw in situ. European Journal of Soil Science 48, 295–300. doi:10.1111/j.1365-2389.1997.tb00549.x

Bach, E.M., Williams, R.J., Hargreaves, S.K., Yang, F., Hofmockel, K.S., 2018. Greatest soil microbial diversity found in micro-habitats. Soil Biology and Biochemistry 118, 217–226. doi:10.1016/j.soilbio.2017.12.018

Bailey, V.L., Pries, C.H., Lajtha, K., 2019. What do we know about soil carbon destabilization? Environmental Research Letters 14, 083004. doi:10.1088/1748-9326/ab2c11

Belnap, J., Eldridge, D., 2001. Biological Soil Crusts: Structure, Function, and Management. Ecological Studies 363–383. doi:10.1007/978-3-642-56475-8_27

Benjamini, Y., Hochberg, Y., 1995. Controlling the False Discovery Rate: A Practical and Powerful Approach to Multiple Testing. Journal of the Royal Statistical Society: Series B (Methodological) 57, 289–300. doi:10.1111/j.2517-6161.1995.tb02031.x

Bergmann, G.T., Bates, S.T., Eilers, K.G., Lauber, C.L., Caporaso, J.G., Walters, W.A., Knight, R., Fierer, N., 2011. The under-recognized dominance of Verrucomicrobia in soil bacterial communities. Soil Biology and Biochemistry 43, 1450–1455. doi:10.1016/j.soilbio.2011.03.012

Bhattacharyya, R., Rabbi, S.M.F., Zhang, Y., Young, I.M., Jones, A.R., Dennis, P.G., Menzies, N.W., Kopittke, P.M., Dalal, R.C., 2021. Soil organic carbon is significantly associated with the pore geometry, microbial diversity and enzyme activity of the macro-aggregates under different land uses. Science of The Total Environment 778, 146286. doi:10.1016/j.scitotenv.2021.146286

Biesgen, D., Frindte, K., Maarastawi, S., Knief, C., 2020. Clay content modulates differences in bacterial community structure in soil aggregates of different size. Geoderma 376, 114544. doi:10.1016/j.geoderma.2020.114544

Bolyen, E., Rideout, J.R., Dillon, M.R., Bokulich, N.A., Abnet, C.C., Al-Ghalith, G.A., Alexander, H., Alm, E.J., Arumugam, M., Asnicar, F., Bai, Y., Bisanz, J.E., Bittinger, K., Brejnrod, A., Brislawn, C.J., Brown, C.T., Callahan, B.J., Caraballo-Rodríguez, A.M., Chase, J., Cope, E.K., Silva, R.D., Diener, C., Dorrestein, P.C., Douglas, G.M., Durall, D.M., Duvallet, C., Edwardson, C.F., Ernst, M., Estaki, M., Fouquier, J., Gauglitz, J.M., Gibbons, S.M., Gibson, D.L., González, A., Gorlick, K., Guo, J., Hillmann, B., Holmes, S., Holste, H., Huttenhower, C., Huttley, G.A., Janssen, S., Jarmusch, A.K., Jiang, L., Kaehler, B.D., Kang, K.B., Keefe, C.R., Keim, P., Kelley, S.T., Knights, D., Koester, I., Kosciolek, T., Kreps, J., Langille, M.G.I., Lee, J., Ley, R., Liu, Y.-X., Loftfield, E., Lozupone, C., Maher, M., Marotz, C., Martin, B.D., McDonald, D., McIver, L.J., Melnik, A.V., Metcalf, J.L., Morgan, S.C., Morton, J.T., Naimey, A.T., Navas-Molina, J.A., Nothias, L.F., Orchanian, S.B., Pearson, T., Peoples, S.L., Petras, D., Preuss, M.L., Pruesse, E., Rasmussen, L.B., Rivers, A., Robeson, M.S., Rosenthal, P., Segata, N., Shaffer, M., Shiffer, A., Sinha, R., Song, S.J., Spear, J.R., Swafford, A.D., Thompson, L.R., Torres, P.J., Trinh, P., Tripathi, A., Turnbaugh, P.J., Ul-Hasan, S., Hooft, J.J.J. van der, Vargas, F., Vázquez-Baeza, Y., Vogtmann, E., Hippel, M. von, Walters, W., Wan, Y., Wang, M., Warren, J., Weber, K.C., Williamson, C.H.D., Willis, A.D., Xu, Z.Z., Zaneveld, J.R., Zhang, Y., Zhu, Q., Knight, R., Caporaso, J.G., 2019. Reproducible, interactive, scalable and extensible microbiome data science using QIIME 2. Nature Biotechnology 37, 852–857. doi:10.1038/s41587-019-0209-9

Braus, M.J., Whitman, T.L., 2021. Standard and non-standard measurements of acidity and the bacterial ecology of northern temperate mineral soils. Soil Biology and Biochemistry 160, 108323. doi:10.1016/j.soilbio.2021.108323

Bray, J.R., Curtis, J.T., 1957. An Ordination of the Upland Forest Communities of Southern Wisconsin. Ecological Monographs 27, 325–349. doi:10.2307/1942268

Caldwell, A.C., Farnham, R.S., Hammers, F.L., 1955. A Chemical and Mineralogical Study of Clay Materials from Several Gray-Brown Podzolic Soils of Minnesota. Soil Science Society of America Journal 19, 351–354. doi:10.2136/sssaj1955.03615995001900030025x

Callahan, B.J., McMurdie, P.J., Rosen, M.J., Han, A.W., Johnson, A.J.A., Holmes, S.P., 2016. DADA2: High-resolution sample inference from Illumina amplicon data. Nature Methods 13, 581–583. doi:10.1038/nmeth.3869

Campbell, C.D., Chapman, S.J., Cameron, C.M., Davidson, M.S., Potts, J.M., 2003. A Rapid Microtiter Plate Method To Measure Carbon Dioxide Evolved from Carbon Substrate Amendments so as To Determine the Physiological Profiles of Soil Microbial Communities by Using Whole Soil. Applied and Environmental Microbiology 69, 3593–3599. doi:10.1128/aem.69.6.3593-3599.2003

Carini, P., Marsden, P.J., Leff, J.W., Morgan, E.E., Strickland, M.S., Fierer, N., 2016. Relic DNA is abundant in soil and obscures estimates of soil microbial diversity. Nature Microbiology 1–6. doi:10.1038/nmicrobiol.2016.242

Cates, A.M., Ruark, M.D., Hedtcke, J.L., Posner, J.L., 2016. Long-term tillage, rotation and perennialization effects on particulate and aggregate soil organic matter. Soil and Tillage Research 155, 371–380. doi:10.1016/j.still.2015.09.008

Chamberlain, L.A., Whitman, T., Ané, J.-M., Diallo, T., Gaska, J.M., Lauer, J.G., Mourtzinis, S., Conley, S.P., 2021. Corn-soybean rotation, tillage, and foliar fungicides: Impacts on yield and soil fungi. Field Crops Research 262, 108030. doi:10.1016/j.fcr.2020.108030

Chase, J.M., Kraft, N.J.B., Smith, K.G., Vellend, M., Inouye, B.D., 2011. Using null models to disentangle variation in community dissimilarity from variation in α-diversity. Ecosphere 2, 1–11. doi:10.1890/es10-00117.1@10.1002/(issn)2150-8925(cat)virtualissue(vi)ecs2

Constancias, F., Prévost-Bouré, N.C., Terrat, S., Aussems, S., Nowak, V., Guillemin, J.-P., Bonnotte, A., Biju-Duval, L., Navel, A., Martins, J.M., Maron, P.-A., Ranjard, L., 2013. Microscale evidence for a high decrease of soil bacterial density and diversity by cropping. Agronomy for Sustainable Development 34, 831–840. doi:10.1007/s13593-013-0204-3

Cruse, R.M., Mier, R., Mize, C.W., 2001. Surface Residue Effects on Erosion of Thawing Soils. Soil Science Society of America Journal 65, 178–184. doi:10.2136/sssaj2001.651178x

Davinic, M., Fultz, L.M., Acosta-Martinez, V., Calderón, F.J., Cox, S.B., Dowd, S.E., Allen, V.G., Zak, J.C., Moore-Kucera, J., 2012. Pyrosequencing and mid-infrared spectroscopy reveal distinct aggregate stratification of soil bacterial communities and organic matter composition. Soil Biology and Biochemistry 46, 63–72. doi:10.1016/j.soilbio.2011.11.012

DeGryze, S., Six, J., Merckx, R., 2006. Quantifying water-stable soil aggregate turnover and its implication for soil organic matter dynamics in a model study. European Journal of Soil Science 57, 693–707. doi:10.1111/j.1365-2389.2005.00760.x

Denef, K., Six, J., Merckx, R., Paustian, K., 2004. Carbon Sequestration in Microaggregates of No-Tillage Soils with Different Clay Mineralogy. Soil Science Society of America Journal 68, 1935–1944. doi:10.2136/sssaj2004.1935

Dini-Andreote, F., Stegen, J.C., Elsas, J.D. van, Salles, J.F., 2015. Disentangling mechanisms that mediate the balance between stochastic and deterministic processes in microbial succession. Proceedings of the National Academy of Sciences of the United States of America 112, E1326–32. doi:10.1073/pnas.1414261112

Dolliver, H., Gupta, S., 2008. Antibiotic Losses in Leaching and Surface Runoff from Manure- Amended Agricultural Land. Journal of Environmental Quality 37, 1227–1237. doi:10.2134/jeq2007.0392

Edwards, A.P., Bremner, J.M., 1967. Microaggregates in Soils. Journal of Soil Science 18, 64– 73. doi:10.1111/j.1365-2389.1967.tb01488.x

Elliott, E.T., 1986. Aggregate Structure and Carbon, Nitrogen, and Phosphorus in Native and Cultivated Soils. Soil Science Society of America Journal 50, 627–633. doi:10.2136/sssaj1986.03615995005000030017x

Faith, D.P., 1992. Conservation evaluation and phylogenetic diversity. Biological Conservation 61, 1–10. doi:10.1016/0006-3207(92)91201-3

Figuerola, E.L.M., Guerrero, L.D., Türkowsky, D., Wall, L.G., Erijman, L., 2015. Reduced turnover of soil bacterial communities. Environmental Microbiology 17, 678–688. doi:10.1111/1462-2920.12497

Frey, S.D., Elliott, E.T., Paustian, K., 1999. Bacterial and fungal abundance and biomass in conventional and no-tillage agroecosystems along two climatic gradients. Soil Biology and Biochemistry 31, 573–585. doi:10.1016/s0038-0717(98)00161-8

Garland, G., Bünemann, E.K., Oberson, A., Frossard, E., Snapp, S., Chikowo, R., Six, J., 2018. Phosphorus cycling within soil aggregate fractions of a highly weathered tropical soil: A conceptual model. Soil Biology and Biochemistry 116, 91–98. doi:10.1016/j.soilbio.2017.10.007

Gupta, S., Munyankusi, E., Moncrief, J., Zvomuya, F., Hanewall, M., 2004. Tillage and Manure Application Effects on Mineral Nitrogen Leaching from Seasonally Frozen Soils. Journal of Environmental Quality 33, 1238–1246. doi:10.2134/jeq2004.1238

Huang, S., Sun, Y.-N., Rui, W.-Y., Liu, W.-R., Zhang, W.-J., 2010. Long-Term Effect of No- Tillage on Soil Organic Carbon Fractions in a Continuous Maize Cropping System of Northeast China. Pedosphere 20, 285–292. doi:10.1016/s1002-0160(10)60016-1

Hutchinson, G.E., 1957. Concluding remarks. Cold Spring Harbor Symposia. Quantitative Biology 22, 415–427.

Janzen, H.H., 2006. The soil carbon dilemma: Shall we hoard it or use it? Soil Biology and Biochemistry 38, 419–424. doi:10.1016/j.soilbio.2005.10.008

Jastrow, J.D., Miller, R.M., 1998. Soil aggregate stabilization and carbon sequestration: Feedbacks through organomineral associations, in: Soil Processes and the Carbon Cycle. pp. 207–223. doi:10.1201/9780203739273-15

Junkins, E.N., McWhirter, J.B., McCall, L.-I., Stevenson, B.S., 2022. Environmental structure impacts microbial composition and secondary metabolism. ISME Communications 2, 15. doi:10.1038/s43705-022-00097-5

Kembel, S.W., Cowan, P.D., Helmus, M.R., Cornwell, W.K., Morlon, H., Ackerly, D.D., Blomberg, S.P., Webb, C.O., 2010. Picante: R tools for integrating phylogenies and ecology. Bioinformatics 26, 1463–1464. doi:10.1093/bioinformatics/btq166

King, A.E., Congreves, K.A., Deen, B., Dunfield, K.E., Voroney, R.P., Wagner-Riddle, C., 2019. Quantifying the relationships between soil fraction mass, fraction carbon, and total soil carbon to assess mechanisms of physical protection. Soil Biology and Biochemistry 135, 95– 107. doi:10.1016/j.soilbio.2019.04.019

Klappenbach, J.A., Dunbar, J.M., Schmidt, T.M., 2000. rRNA operon copy number reflects ecological strategies of bacteria. Appl. Environ. Microbiol. 66, 1328–1333. doi:10.1128/aem.66.4.1328-1333.2000

Kozich, J.J., Westcott, S.L., Baxter, N.T., Highlander, S.K., Schloss, P.D., 2013. Development of a Dual-Index Sequencing Strategy and Curation Pipeline for Analyzing Amplicon Sequence Data on the MiSeq Illumina Sequencing Platform. Appl. Environ. Microbiol. 79, 5112–5120. doi:10.1128/aem.01043-13

Kravchenko, A.N., Guber, A.K., Razavi, B.S., Koestel, J., Quigley, M.Y., Robertson, G.P., Kuzyakov, Y., 2019. Microbial spatial footprint as a driver of soil carbon stabilization. Nature Communications 10, 3121. doi:10.1038/s41467-019-11057-4

Kuzyakov, Y., Blagodatskaya, E., 2015. Microbial hotspots and hot moments in soil: Concept & review. Soil Biology and Biochemistry 83, 184–199. doi:10.1016/j.soilbio.2015.01.025

Legendre, P., Gallagher, E.D., 2001. Ecologically meaningful transformations for ordination of species data. Oecologia 129, 271–280. doi:10.1007/s004420100716

LeGuillou, C., Angers, D.A., Leterme, P., Menasseri-Aubry, S., 2012. Changes during winter in water-stable aggregation due to crop residue quality. Soil Use and Management 28, 590–595. doi:10.1111/j.1475-2743.2012.00427.x

Lehmann, A., Rillig, M.C., 2015. Understanding mechanisms of soil biota involvement in soil aggregation: A way forward with saprobic fungi? Soil Biology and Biochemistry 88, 298– 302. doi:10.1016/j.soilbio.2015.06.006

Lennon, J.T., Jones, S.E., 2011. Microbial seed banks: the ecological and evolutionary implications of dormancy. Nature Publishing Group 9, 119–130. doi:10.1038/nrmicro2504

Liu, Y.J., Laird, D.A., Barak, P., 1997. Release and Fixation of Ammonium and Potassium under Long-Term Fertility Management. Soil Science Society of America Journal 61, 310–314. doi:10.2136/sssaj1997.03615995006100010044x

Martin, B.D., Witten, D., Willis, A.D., 2021. corncob: Count Regression for Correlated Observations with the Beta-Binomial.

McMurdie, P.J., Holmes, S., 2013. phyloseq: An R Package for Reproducible Interactive Analysis and Graphics of Microbiome Census Data. PLOS ONE 8, e61217. doi:10.1371/journal.pone.0061217

Nemergut, D.R., Knelman, J.E., Ferrenberg, S., Bilinski, T., Melbourne, B., Jiang, L., Violle, C., Darcy, J.L., Prest, T., Schmidt, S.K., Townsend, A.R., 2016. Decreases in average bacterial community rRNA operon copy number during succession. The ISME Journal 10, 1147–1156. doi:10.1038/ismej.2015.191

Ning, D., Yuan, M., Wu, L., Zhang, Y., Guo, X., Zhou, X., Yang, Y., Arkin, A.P., Firestone, M.K., Zhou, J., 2020. A quantitative framework reveals ecological drivers of grassland microbial community assembly in response to warming. Nature Communications 11, 4717. doi:10.1038/s41467-020-18560-z

Oades, J., Waters, A., 1991. Aggregate hierarchy in soils. Soil Research 29, 815–828. doi:10.1071/sr9910815

Oades, J.M., 1984. Soil organic matter and structural stability: mechanisms and implications for management. Plant and Soil 76, 319–337. doi:10.1007/bf02205590

Ogle, S.M., Alsaker, C., Baldock, J., Bernoux, M., Breidt, F.J., McConkey, B., Regina, K., Vazquez-Amabile, G.G., 2019. Climate and Soil Characteristics Determine Where No-Till Management Can Store Carbon in Soils and Mitigate Greenhouse Gas Emissions. Scientific Reports 9, 11665. doi:10.1038/s41598-019-47861-7

Or, D., Keller, T., Schlesinger, W.H., 2021. Natural and managed soil structure: On the fragile scaffolding for soil functioning. Soil and Tillage Research 208, 104912. doi:10.1016/j.still.2020.104912

Ordóñez, R.A., Archontoulis, S.V., Martinez-Feria, R., Hatfield, J.L., Wright, E.E., Castellano, M.J., 2020. Root to shoot and carbon to nitrogen ratios of maize and soybean crops in the US Midwest. European Journal of Agronomy 120, 126130. doi:10.1016/j.eja.2020.126130

Paustian, K., Collins, H.P., Paul, E.A., 1997. Management Controls on Soil Carbon, in: Soil Organic Matter in Temperate Agroecosystems. CRC Press, Inc., pp. 15–49. doi:10.1201/9780367811693-2

Paustian, K., Six, J., Elliott, E.T., Hunt, H.W., 2000. Management options for reducing CO2 emissions from agricultural soils. Biogeochemistry 48, 147–163. doi:10.1023/a:1006271331703

Pedersen, P., Lauer, J.G., 2003. Corn and Soybean Response to Rotation Sequence, Row Spacing, and Tillage System. Agronomy Journal 95, 965–971. doi:10.2134/agronj2003.9650

Pérez-Valera, E., Goberna, M., Verdú, M., 2015. Phylogenetic structure of soil bacterial communities predicts ecosystem functioning. FEMS Microbiology Ecology 91, fiv031. doi:10.1093/femsec/fiv031

Perfect, E., Kay, B.D., Loon, W.K.P., Sheard, R.W., Pojasok, T., 1990. Factors Influencing Soil Structural Stability within a Growing Season. Soil Science Society of America Journal 54, 173–179. doi:10.2136/sssaj1990.03615995005400010027x

Piazza, G., Pellegrino, E., Moscatelli, M.C., Ercoli, L., 2020. Long-term conservation tillage and nitrogen fertilization effects on soil aggregate distribution, nutrient stocks and enzymatic activities in bulk soil and occluded microaggregates. Soil and Tillage Research 196, 104482. doi:10.1016/j.still.2019.104482

Powlson, D.S., Stirling, C.M., Jat, M.L., Gerard, B.G., Palm, C.A., Sanchez, P.A., Cassman, K.G., 2014. Limited potential of no-till agriculture for climate change mitigation. Nature Climate Change 4, 678–683. doi:10.1038/nclimate2292

Quast, C., Pruesse, E., Yilmaz, P., Gerken, J., Schweer, T., Yarza, P., Peplies, J., Glöckner, F.O., 2013. The SILVA ribosomal RNA gene database project: improved data processing and web- based tools. Nucleic Acids Research 41, D590–D596. doi:10.1093/nar/gks1219

Ranjard, L., Poly, F., Combrisson, J., Richaume, A., Gourbière, F., Thioulouse, J., Nazaret, S., 2000. Heterogeneous Cell Density and Genetic Structure of Bacterial Pools Associated with Various Soil Microenvironments as Determined by Enumeration and DNA Fingerprinting Approach (RISA). Microbial Ecology 39, 263–272. doi:10.1007/s002480000032

Ranjard, L., Richaume, A., 2001. Quantitative and qualitative microscale distribution of bacteria in soil. Research in Microbiology 152, 707–716. doi:10.1016/s0923-2508(01)01251-7

R-Core-Team, 2018. R: A Language and Environment for Statistical Computing, R Foundation for Statistical Computing.

Rillig, M.C., Muller, L.A., Lehmann, A., 2017. Soil aggregates as massively concurrent evolutionary incubators. The ISME Journal 11, 1943–1948. doi:10.1038/ismej.2017.56

Roller, B.R.K., Stoddard, S.F., Schmidt, T.M., 2016. Exploiting rRNA Operon Copy Number to Investigate Bacterial Reproductive Strategies. Nature Microbiology 1, 16160–16160. doi:10.1038/nmicrobiol.2016.160

Sae-Tun, O., Bodner, G., Rosinger, C., Zechmeister-Boltenstern, S., Mentler, A., Keiblinger, K., 2022. Fungal biomass and microbial necromass facilitate soil carbon sequestration and aggregate stability under different soil tillage intensities. Applied Soil Ecology 179, 104599. doi:10.1016/j.apsoil.2022.104599

Santoni, M., Verdi, L., Pathan, S.I., Napoli, M., Marta, A.D., Dani, F.R., Pacini, G.C., Ceccherini, M.T., 2022. Soil microbiome biomass, activity, composition and CO2 emissions in a long-term organic and conventional farming systems. Soil Use and Management. doi:10.1111/sum.12836

Sarkar, B., Singh, M., Mandal, S., Churchman, G.J., Bolan, N.S., 2018. The Future of Soil Carbon. pp. 71–86. doi:10.1016/b978-0-12-811687-6.00003-1

Schimel, J.P., Schaeffer, S.M., 2012. Microbial control over carbon cycling in soil. Frontiers in Microbiology 3, 348–11. doi:10.3389/fmicb.2012.00348

Schmidt, R., Gravuer, K., Bossange, A.V., Mitchell, J., Scow, K., 2018. Long-term use of cover crops and no-till shift soil microbial community life strategies in agricultural soil. PLOS ONE 13, e0192953. doi:10.1371/journal.pone.0192953

Sexstone, A.J., Revsbech, N.P., Parkin, T.B., Tiedje, J.M., 1985. Direct Measurement of Oxygen Profiles and Denitrification Rates in Soil Aggregates. Soil Science Society of America Journal 49, 645–651. doi:10.2136/sssaj1985.03615995004900030024x

Sheehy, J., Regina, K., Alakukku, L., Six, J., 2015. Impact of no-till and reduced tillage on aggregation and aggregate-associated carbon in Northern European agroecosystems. Soil and Tillage Research 150, 107–113. doi:10.1016/j.still.2015.01.015

Simpson, R.T., Frey, S.D., Six, J., Thiet, R.K., 2004. Preferential Accumulation of Microbial Carbon in Aggregate Structures of No-Tillage Soils. Soil Science Society of America Journal 68, 1249–1255. doi:10.2136/sssaj2004.1249

Six, J., Bossuyt, H., Degryze, S., Denef, K., 2004. A history of research on the link between (micro)aggregates, soil biota, and soil organic matter dynamics. Soil and Tillage Research 79, 7–31. doi:10.1016/j.still.2004.03.008

Six, J., Callewaert, P., Lenders, S., Gryze, S.D., Morris, S.J., Gregorich, E.G., Paul, E.A., Paustian, K., 2002. Measuring and Understanding Carbon Storage in Afforested Soils by Physical Fractionation. Soil Science Society of America Journal 66, 1981–1987. doi:10.2136/sssaj2002.1981

Six, J., Elliott, E.T., Paustian, K., 2000a. Soil macroaggregate turnover and microaggregate formation: a mechanism for C sequestration under no-tillage agriculture. Soil Biology and Biochemistry 32, 2099–2103. doi:10.1016/s0038-0717(00)00179-6

Six, J., Elliott, E.T., Paustian, K., 1999. Aggregate and Soil Organic Matter Dynamics under Conventional and No-Tillage Systems. Soil Science Society of America Journal 63, 1350– 1358. doi:10.2136/sssaj1999.6351350x

Six, J., Elliott, E.T., Paustian, K., Doran, J.W., 1998. Aggregation and Soil Organic Matter Accumulation in Cultivated and Native Grassland Soils. Soil Science Society of America Journal 62, 1367–1377. doi:10.2136/sssaj1998.03615995006200050032x

Six, J., Frey, S.D., Thiet, R.K., Batten, K.M., 2006. Bacterial and Fungal Contributions to Carbon Sequestration in Agroecosystems. Soil Science Society of America Journal 70, 555. doi:10.2136/sssaj2004.0347

Six, J., Paustian, K., 2014. Aggregate-associated soil organic matter as an ecosystem property and a measurement tool. Soil Biology and Biochemistry 68, A4–A9. doi:10.1016/j.soilbio.2013.06.014

Six, J., Paustian, K., Elliott, E.T., Combrink, C., 2000b. Soil Structure and Organic Matter I. Distribution of Aggregate-Size Classes and Aggregate-Associated Carbon. Soil Science Society of America Journal 64, 681–689. doi:10.2136/sssaj2000.642681x

Smith, C.R., Blair, P.L., Boyd, C., Cody, B., Hazel, A., Hedrick, A., Kathuria, H., Khurana, P., Kramer, B., Muterspaw, K., Peck, C., Sells, E., Skinner, J., Tegeler, C., Wolfe, Z., 2016. Microbial community responses to soil tillage and crop rotation in a corn/soybean agroecosystem. Ecology and Evolution 6, 8075–8084. doi:10.1002/ece3.2553

Srour, A.Y., Ammar, H.A., Subedi, A., Pimentel, M., Cook, R.L., Bond, J., Fakhoury, A.M., 2020. Microbial Communities Associated With Long-Term Tillage and Fertility Treatments in a Corn-Soybean Cropping System. Frontiers in Microbiology 11, 1363. doi:10.3389/fmicb.2020.01363

Stegen, J.C., Lin, X., Fredrickson, J.K., Chen, X., Kennedy, D.W., Murray, C.J., Rockhold, M.L., Konopka, A., 2013. Quantifying community assembly processes and identifying features that impose them. The ISME Journal 7, 2069–2079. doi:10.1038/ismej.2013.93

Stegen, J.C., Lin, X., Fredrickson, J.K., Konopka, A.E., 2015. Estimating and mapping ecological processes influencing microbial community assembly. Frontiers in Microbiology 6. doi:10.3389/fmicb.2015.00370

Stegen, J.C., Lin, X., Konopka, A.E., Fredrickson, J.K., 2012. Stochastic and deterministic assembly processes in subsurface microbial communities. The ISME Journal 6, 1653–1664. doi:10.1038/ismej.2012.22

Stoddard, S.F., Smith, B.J., Hein, R., Roller, B.R.K., Schmidt, T.M., 2015. rrnDB: improved tools for interpreting rRNA gene abundance in bacteria and archaea and a new foundation for future development. Nucleic Acids Research 43, D593–D598. doi:10.1093/nar/gku1201

Tisdall, J.M., Oades, J.M., 1982. Organic matter and water-stable aggregates in soils 33, 141– 163. doi:10.1111/j.1365-2389.1982.tb01755.x

Totsche, K.U., Amelung, W., Gerzabek, M.H., Guggenberger, G., Klumpp, E., Knief, C., Lehndorff, E., Mikutta, R., Peth, S., Prechtel, A., Ray, N., Kögel-Knabner, I., 2018. Microaggregates in soils. Journal of Plant Nutrition and Soil Science 181, 104–136. doi:10.1002/jpln.201600451

Trivedi, P., Delgado-Baquerizo, M., Jeffries, T.C., Trivedi, C., Anderson, I.C., Lai, K., McNee, M., Flower, K., Singh, B.P., Minkey, D., Singh, B.K., 2017. Soil aggregation and associated microbial communities modify the impact of agricultural management on carbon content. Environmental Microbiology 19, 3070–3086. doi:10.1111/1462-2920.13779

Upton, R.N., Bach, E.M., Hofmockel, K.S., 2019. Spatio-temporal microbial community dynamics within soil aggregates. Soil Biology and Biochemistry 132, 58–68. doi:10.1016/j.soilbio.2019.01.016

Vellend, M., 2010. Conceptual Synthesis in Community Ecology. The Quarterly Review of Biology 85, 183–206. doi:10.1086/652373

Vogel, H., Balseiro-Romero, M., Kravchenko, A., Otten, W., Pot, V., Schlüter, S., Weller, U., Baveye, P.C., 2022. A holistic perspective on soil architecture is needed as a key to soil functions. European Journal of Soil Science 73. doi:10.1111/ejss.13152

Walters, W., Hyde, E.R., Berg-Lyons, D., Ackermann, G., Humphrey, G., Parada, A., Gilbert, J.A., Jansson, J.K., Caporaso, J.G., Fuhrman, J.A., Apprill, A., Knight, R., Bik, H., 2016. Improved Bacterial 16S rRNA Gene (V4 and V4-5) and Fungal Internal Transcribed Spacer Marker Gene Primers for Microbial Community Surveys. MSystems 1, e00009–15. doi:10.1128/msystems.00009-15

West, J.R., Whitman, T., 2022. Disturbance by soil mixing decreases microbial richness and supports homogenizing community assembly processes. FEMS Microbiology Ecology. doi:10.1093/femsec/fiac089

Whitman, T.L., Whitman, E., Woolet, J., Flannigan, M.D., Thompson, D.K., Parisien, M.-A., 2019. Soil bacterial and fungal response to wildfires in the Canadian boreal forest across a burn severity gradient. Soil Biology and Biochemistry 107571–59. doi:10.1016/j.soilbio.2019.107571

Wickham, H., 2016. ggplot2: Elegant Graphics for Data Analysis, Springer-Verlag.

Wilhelm, R.C., Amsili, J.P., Kurtz, K.S.M., Es, H.M. van, Buckley, D.H., 2023. Ecological insights into soil health according to the genomic traits and environment-wide associations of bacteria in agricultural soils. ISME Communications 3, 1. doi:10.1038/s43705-022-00209-1

Willis, A.D., Bunge, J., Whitman, T.L., 2017. Improved detection of changes in species richness in high diversity microbial communities. Journal of the Royal Statistical Society: Series C (Applied Statistics) 66, 963–977. doi:10.1111/rssc.12206

Wilpiszeski, R.L., Aufrecht, J.A., Retterer, S.T., Sullivan, M.B., Graham, D.E., Pierce, E.M., Zablocki, O.D., Palumbo, A.V., Elias, D.A., 2019. Soil Aggregate Microbial Communities: Towards Understanding Microbiome Interactions at Biologically Relevant Scales. Applied and Environmental Microbiology 85, 689. doi:10.1128/aem.00324-19

Woolf, D., Lehmann, J., 2019. Microbial models with minimal mineral protection can explain long-term soil organic carbon persistence. Scientific Reports 9, 6522. doi:10.1038/s41598-019-43026-8

Yamada, T., Sekiguchi, Y., 2020. Bergey’s Manual of Systematics of Archaea and Bacteria 1–2. doi:10.1002/9781118960608.cbm00064

Yilmaz, P., Parfrey, L.W., Yarza, P., Gerken, J., Pruesse, E., Quast, C., Schweer, T., Peplies, J., Ludwig, W., Glöckner, F.O., 2013. The SILVA and “All-species Living Tree Project (LTP)” taxonomic frameworks. Nucleic Acids Research 42, D643–D648. doi:10.1093/nar/gkt1209

Zheng, H., Liu, W., Zheng, J., Luo, Y., Li, R., Wang, H., Qi, H., 2018. Effect of long-term tillage on soil aggregates and aggregate-associated carbon in black soil of Northeast China. PLOS ONE 13, e0199523. doi:10.1371/journal.pone.0199523

Zhou, J., Ning, D., 2017. Stochastic Community Assembly: Does It Matter in Microbial Ecology? Microbiology and Molecular Biology Reviews : MMBR 81, 1–32. doi:10.1128/mmbr.00002-17

Zuber, S.M., Villamil, M.B., 2016. Meta-analysis approach to assess effect of tillage on microbial biomass and enzyme activities. Soil Biology and Biochemistry 97, 176–187. doi:10.1016/j.soilbio.2016.03.011

