## Supplementary_Information for "Tillage homogenizes soil bacterial communities in microaggregate fractions by facilitating dispersal"

#### Method: PCR amplification, and 16S rRNA library prep

All DNA was stored at or below −20 °C throughout the experiment. Each round of soil extraction included one extraction blank, which was carried through the following pipeline alongside soil samples. 16S rRNA genes of extracted DNA were amplified in triplicate using PCR. Variable region V4 of the 16S rRNA gene was targeted using forward primer 515f and reverse primer 806r (Walters et al., 2016). Primers also contained barcodes and Illumina sequencing adapters (Kozich et al., 2013). The following reagents comprised each 25 µL PCR reaction: 12.5 µL Q5 Hot Start High-Fidelity 2X Master mix (Catalog No. M0494, New England BioLabs, Ipswich, MA), 1.25 µL 515f forward primer (10 mM), 1.25 µL 806r reverse primer (10 mM), 1 µL DNA extract, 1.25 µL Bovine Serum Albumin (20 mg/mL; Catalog No. 97064-342, VWR International, Radnor, PA, and 7.75 µL PCR-grade water. The plate was sealed and briefly centrifuged prior to 30 PCR cycles on an Eppendorf Mastercycler nexus gradient thermal cycler (Hamburg, Germany) using the following parameters: 98 °C for 2 min + 30 × (98 °C for 10 seconds + 58 °C for 15 seconds + 72 °C for 10 seconds) + 72 °C for 2 min and 4 °C hold.

Successful amplification was verified via gel electrophoresis using 1% TAE agarose gel and Invitrogen SYBR Safe DNA Gel Stain (Catalog No. S33102, ThermoFisher Scientific, Carlsbad, CA). Each well was loaded with 5 µL PCR product mixed with 1 µL Purple 6X Gel Loading Dye (Catalog No. B7025S, New England BioLabs, Ipswich, MA). Successful amplification of target base pair length was confirmed using a 2-Log DNA Ladder (0.1–10.0 kilobases; Catalog No. N3200, New England BioLabs, Ipswich, MA) in each gel. Gel ran for ninety minutes at 115V. Gels were photographed and checked visually for amplification.

The SequalPrep Normalization Plate Kit (Catalog No. A1051001, ThermoFisher Scientific, Carlsbad, CA) was used to normalize amplicon yields across samples using a limited binding capacity solid phase. Normalization was performed following kit instructions using 25 μL of the pooled triplicate PCR product, and yielded 20 μL of eluted DNA per sample, which was pooled following elution. The combined DNA library was concentrated using a SpeedVac Vacuum Concentrator System (ThermoFisher Scientific, Waltham, MA) prior to further DNA purification using Wizard SV Gel and PCR Clean-Up System (Catalog No. A9281, Promega Corporation, Madison, WI). Kit instructions were followed except the nuclease-free water was divided into 30 μL and 20 μL increments with the incubation and centrifuge steps after both additions.

#### Supplementary Table

**Table S1**. Soil properties based on representative subsamples for each treatment at each site, 0-5 cm. OM = organic matter; CEC = cation exchange capacity.
Soil texture (percent sand, silt, and clay) was determined via hydrometer method (Bouyoucos, 1962); organic matter was determined via loss on ignition (Schulte and Hopkins, 1996); pH was determined for 1:1 water (Richards, 1954); plant-available P and K were determined by Bray-1 method (Bray and Kurtz, 1945); and plant-available Ca and Mg were determined by ammonium acetate method (Thomas, 1982).

| Site | Treatment | Sand | Silt | Clay | OM | Texture | pH | P | K | Ca | Mg | CEC |
|---|---|---|---|---|---|---|---|---|---|---|---|---|
| | | ---------------- % --------------- | | | | | | ------------- *mg kg$^{-1}$* ------------ | | | | |
| **Arlington** | **No-tillage** | 14 | 64 | 22 | 5.2 | Silt loam | 7 | 50 | 266 | 1890 | 556 | 17 |
| | **Tillage** | 14 | 62 | 24 | 3.8 | Silt loam | 7 | 49 | 247 | 2058 | 599 | 17 |
| **Lancaster** | **No-tillage** | 18 | 64 | 18 | 3.5 | Silt loam | 6.9 | 60 | 171 | 1270 | 393 | 11 |
| | **Tillage** | 8 | 72 | 20 | 2.6 | Silt loam | 7 | 75 | 204 | 1242 | 393 | 10 |

**Table S2**. Taxa enriched in tillage and no-tillage treatments.
*See accompanying file SI_Table_S2_enriched_taxa_by_tillage_trt.csv*

**Table S3**. Taxa enriched in free and occluded microaggregate fractions.
*See accompanying file SI_Table_S3_enriched_taxa_in_microaggregates.csv*

### Supplementary Figures

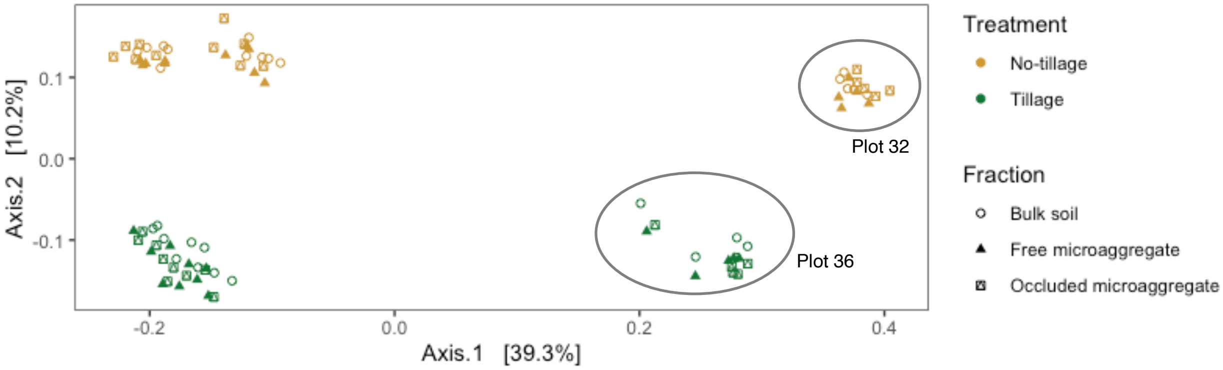

**Figure S1**. Principal coordinates analysis of Bray-Curtis dissimilarities of Hellinger-transformed community relative abundances, colored by mixing treatment for Lancaster, WI, including the two plots (32 and 36) that were excluded from analysis in the main text.
Each point represents the community of one sample-fraction. Soil fractions are as follows: Bulk soil = whole soil; Free microaggregate = 53–250 µm fraction from bulk soil; Occluded microaggregate = 53–250 µm fraction occluded within macroaggregate fraction.

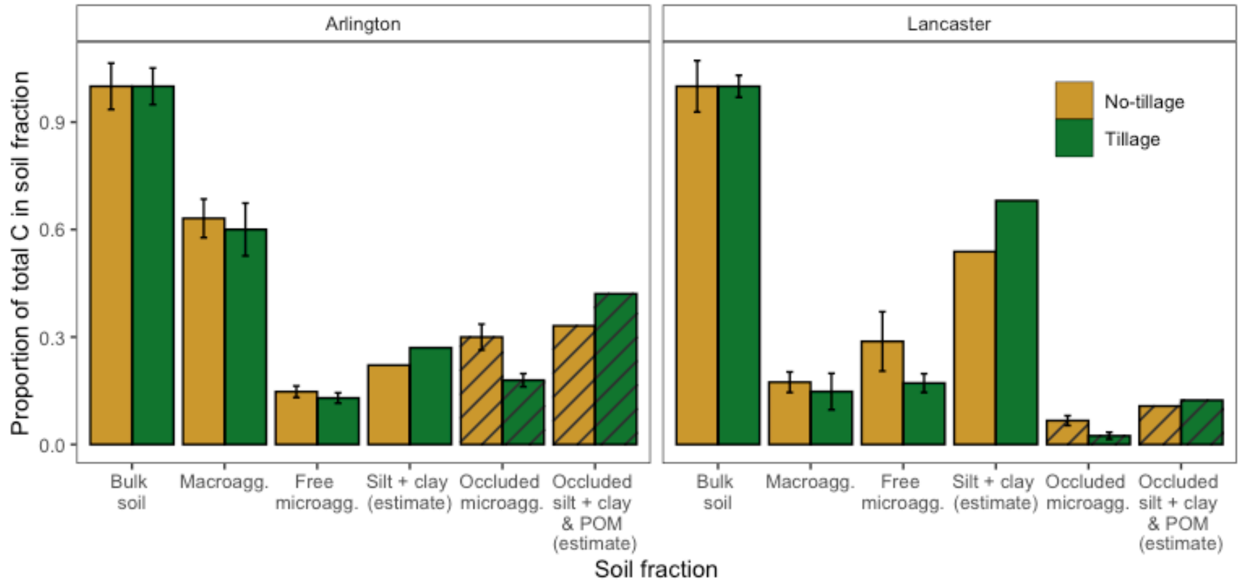

**Figure S2**. Proportion of bulk soil's total C in each soil fraction (i.e., bulk soil = 1.0). Soil fractions are as follows: Bulk soil = whole soil; Macroagg. = macroaggregate fraction, 250–2000 µm; Free microagg. = microaggregate fraction from bulk soil, 53–250 µm; Silt + clay (estimate) = Carbon content in the < 53 µm fraction, estimated to be Bulk soil – (Macroagg. + Free microagg.); Occluded microagg. = microaggregate fraction occluded within macroaggregate fraction, 53–250 µm; Occluded silt + clay & POM (estimate) = Carbon content in the < 53 µm fraction occluded within the macroaggregate fraction, estimated to be Macroagg. – Occluded microagg. Error bars represent ± 1.96 SE. The estimated silt + clay carbon contents do not have associated error bars. Striped bars represent occluded fractions.

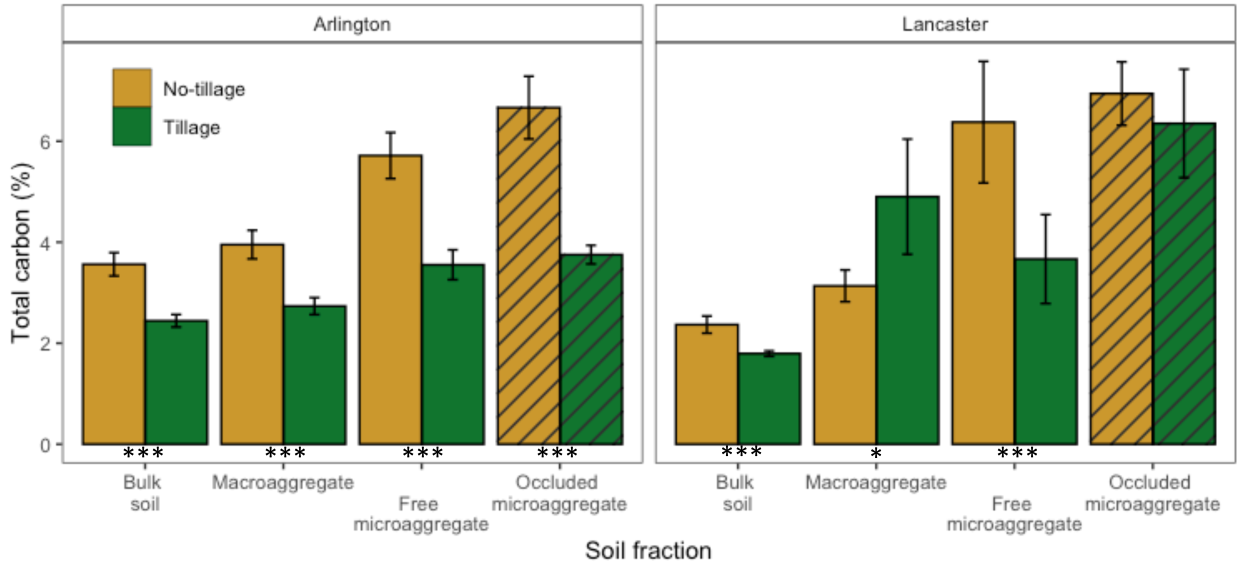

**Figure S3**. Carbon concentration (total C, %), by soil fraction and tillage treatment. Soil fractions are as follows: Bulk soil = sieved field-moist soil < 2000 µm; Macroaggregate = macroaggregate fraction, 250–2000 µm; Free microaggregate = microaggregate fraction from bulk soil, 53–250 µm; Occluded microaggregate = microaggregate fraction occluded within macroaggregate fraction, 53–250 µm. Error bars represent ± 1.96 SE. Asterisks indicate significant tillage treatment differences within soil fraction: *** = $p < 0.001$, ** = $p < 0.01$, * = $p < 0.05$. Striped bars represent occluded fractions.

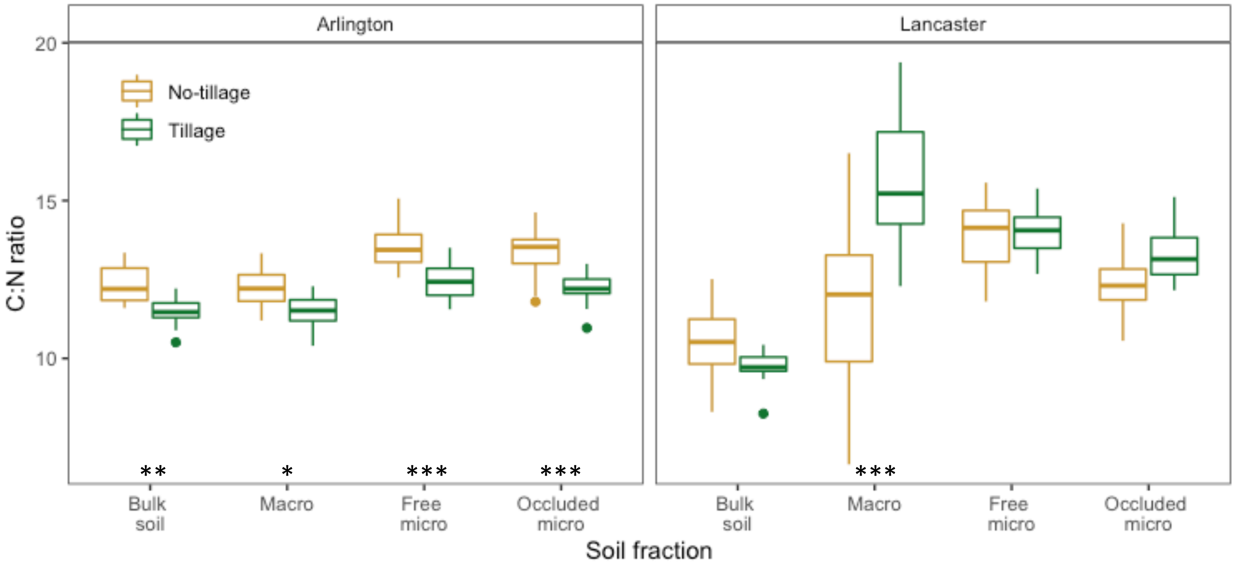

**Figure S4**. C:N ratio, by soil fraction and tillage treatment. Soil fractions are as follows: Bulk soil = whole soil; Macro = macroaggregate fraction, 250–2000 µm; Free micro = microaggregate fraction from bulk soil, 53–250 µm; Occluded micro = microaggregate fraction occluded within macroaggregate fraction, 53–250 µm. Asterisks indicate significant tillage treatment differences, within soil fraction: *** = $p < 0.001$, ** = $p < 0.01$, * = $p < 0.05$.

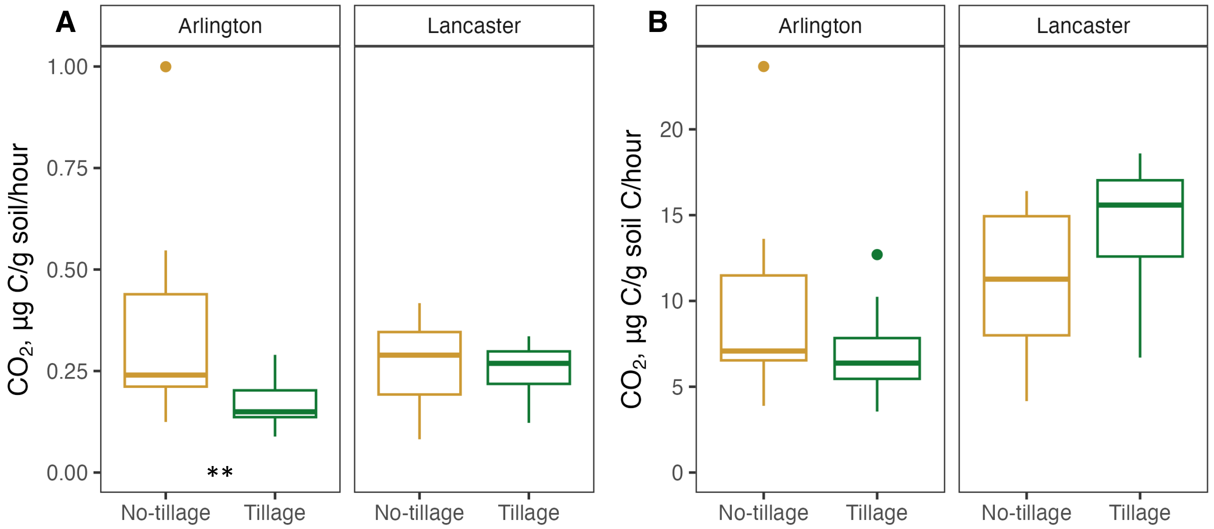

**Figure S5**. Soil respiration, by tillage treatment on a per g soil basis (**A**) and on a per g soil C basis (**B**). $CO_2$ evolution was measured on field moist bulk soil sieved at 2 mm using the MicroResp system (Campbell et al., 2003), a setup designed to capture $CO_2$ from small soil samples using a colorimetric indicator mounted on top of a 96-deep well plate. Asterisks indicate significant tillage treatment differences, within site: *** = $p < 0.001$, ** = $p < 0.01$, * = $p < 0.05$.

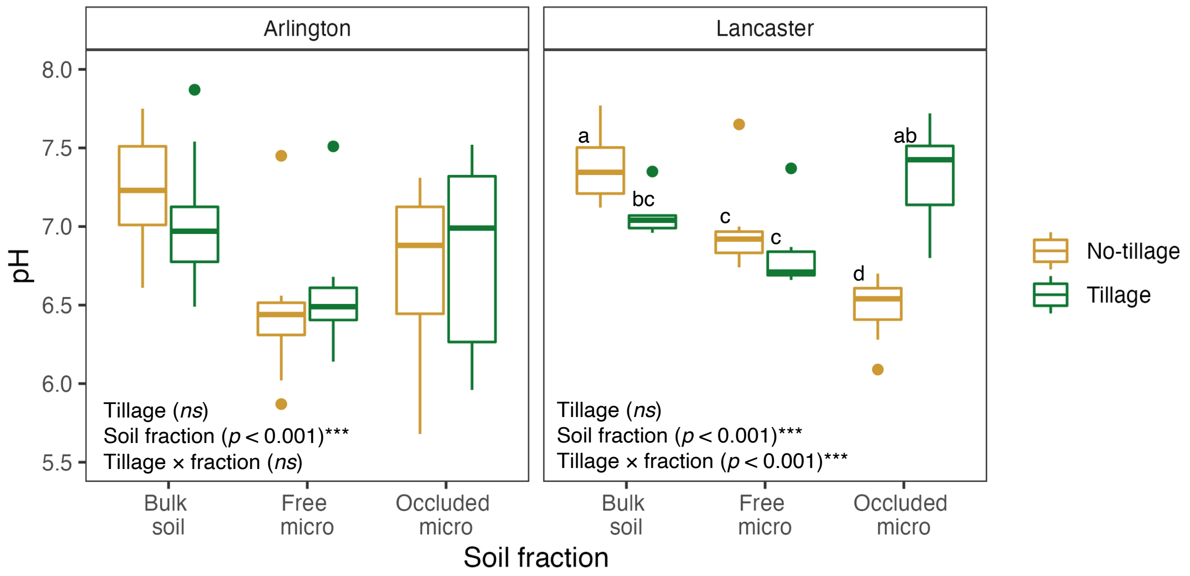

**Figure S6**. Soil pH, by tillage treatment and fraction. Compact letter display, in which boxplots with the same letter are not significantly different, indicates the significant interaction of tillage treatment × soil fraction at the Lancaster site. Bulk soil = whole soil; Free micro = microaggregate fraction from bulk soil, 53–250 µm; Occluded micro = microaggregate fraction occluded within macroaggregate fraction, 53–250 µm.

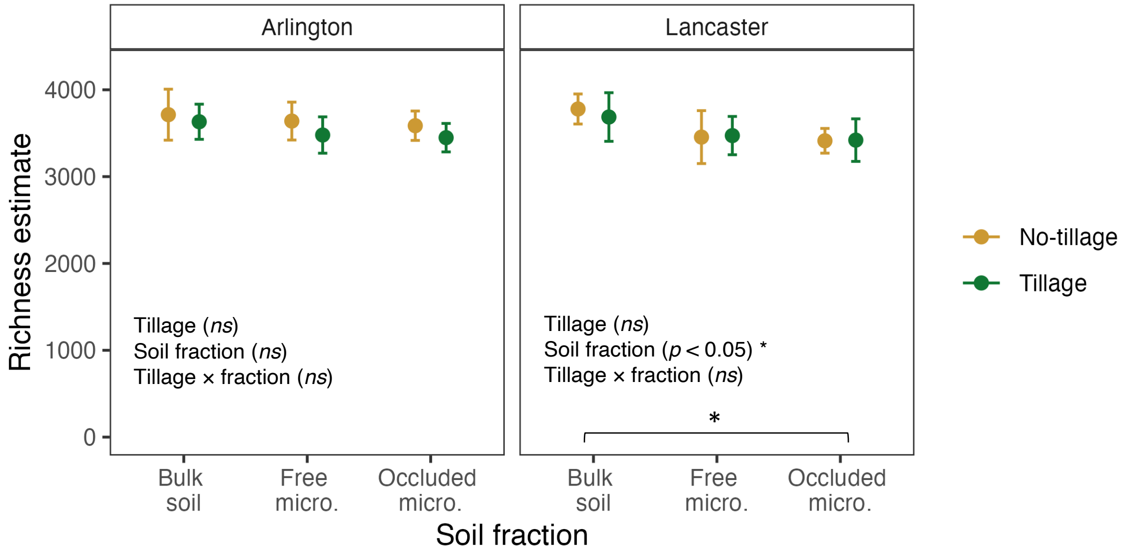

**Figure S7**. OTU richness, by tillage treatment and soil fraction. Richness was estimated using the weighted linear regression model of OTU richness estimates, which weights observations based on variance, using *breakaway::betta* (Willis et al., 2017). Bulk soil = whole soil; Free micro. = microaggregate fraction from bulk soil, 53–250 µm; Occluded micro. = microaggregate fraction occluded within macroaggregate fraction, 53–250 µm. Error bars represent ± 1.96 SE. Asterisk indicates a significant difference between soil fractions: * = $p < 0.05$.

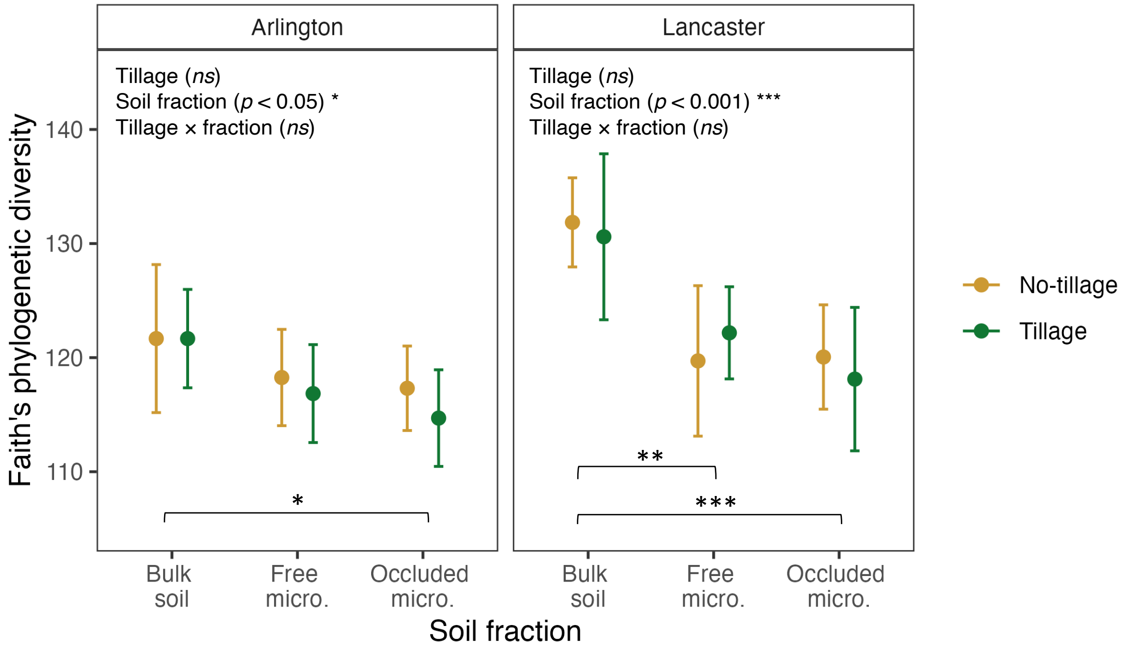

**Figure S8**. Faith's phylogenetic diversity, by tillage treatment and soil fraction. Bulk soil = whole soil; Free micro. = microaggregate fraction from bulk soil, 53–250 µm; Occluded micro. = microaggregate fraction occluded within macroaggregate fraction, 53–250 µm. Error bars represent ± 1.96 SE. Asterisks indicate significant differences between soil fractions: *** = $p < 0.001$, ** = $p < 0.01$, * = $p < 0.05$.

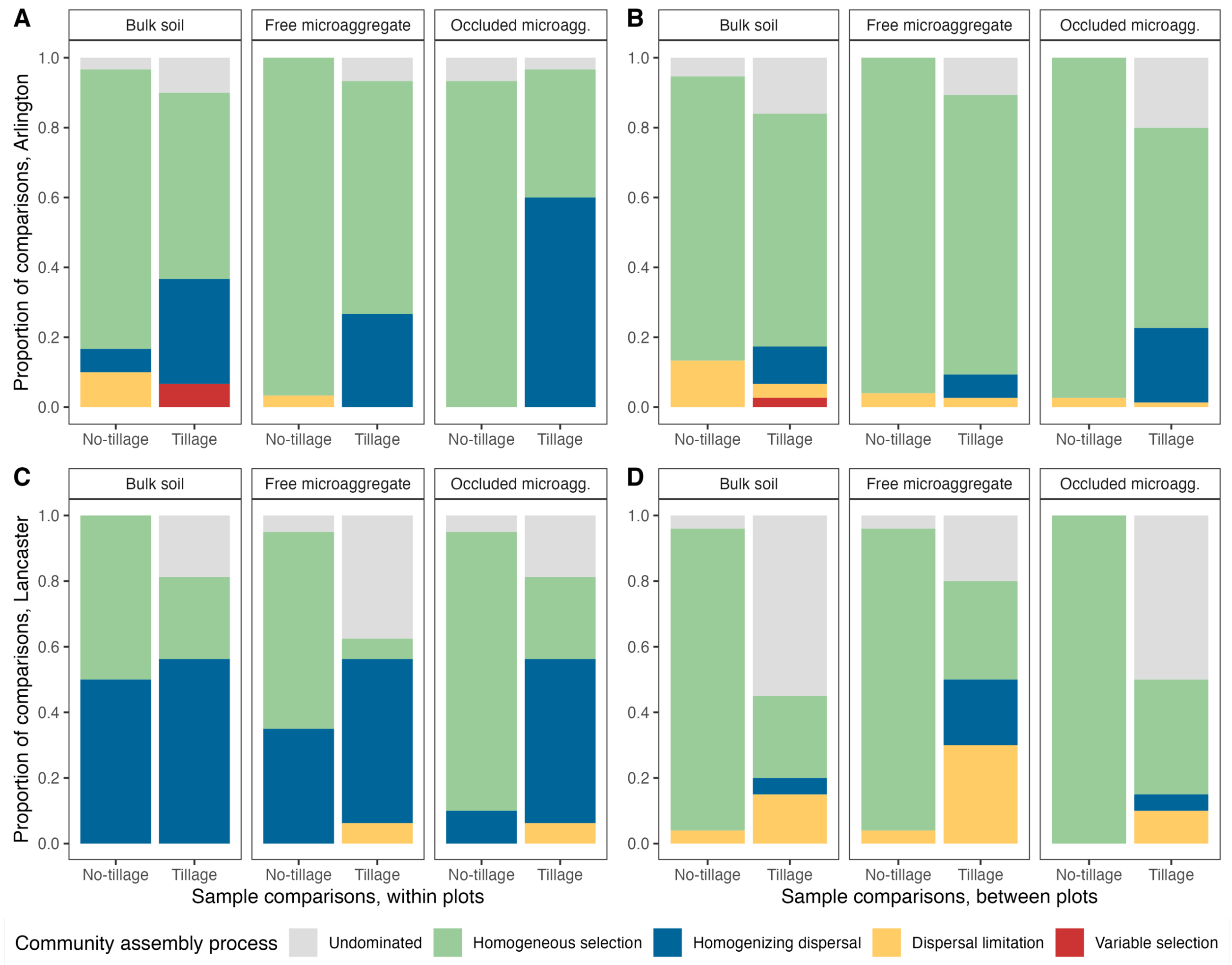

**Figure S9**. Community assembly processes at the full sample community scale, by tillage treatment, within bulk soil, free microaggregate, and occluded microaggregate fractions at Arlington, WI (A & B); and Lancaster, WI (C & D). Sample comparisons were made within plots (A & C) or between plots (B & D). Community assembly processes were estimated using a null-modeling approach within each site × treatment × fraction set of samples, based on sample comparisons for each possible pair of samples, based on the method of Stegen et al. (2012, 2013, 2015). As detailed in the main text, first the influence of selection was determined using the $\beta$-mean nearest taxon distance, and then the influence of dispersal was determined using the modified Raup-Crick metric based on Bray-Curtis dissimilarity. Bulk soil = whole soil; Free microaggregate = microaggregate fraction from bulk soil, 53–250 µm; Occluded microagg. = microaggregate fraction occluded within macroaggregate fraction, 53–250 µm.

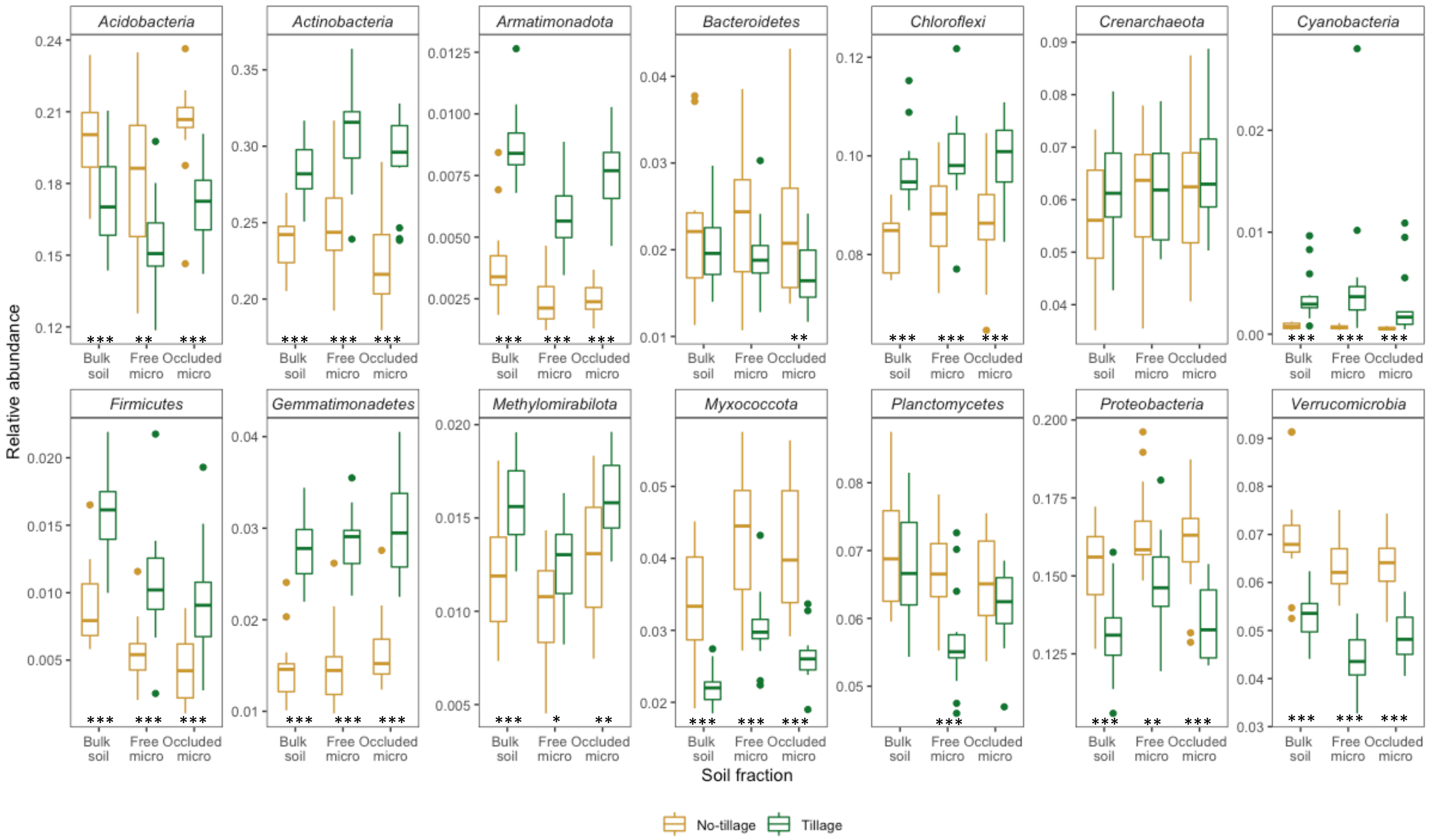

**Figure S10**. Relative abundances of representative phyla at Arlington, WI. Bulk soil = whole soil; Free micro = microaggregate fraction from bulk soil, 53–250 µm; Occluded micro = microaggregate fraction occluded within macroaggregate fraction, 53–250 µm. Asterisks indicate significant treatment differences in relative abundances between tillage treatment, within fraction, determined using *corncob::differentialTest*, with taxa agglomerated at the phylum level: *** = $p < 0.001$, ** = $p < 0.01$, * = $p < 0.05$.

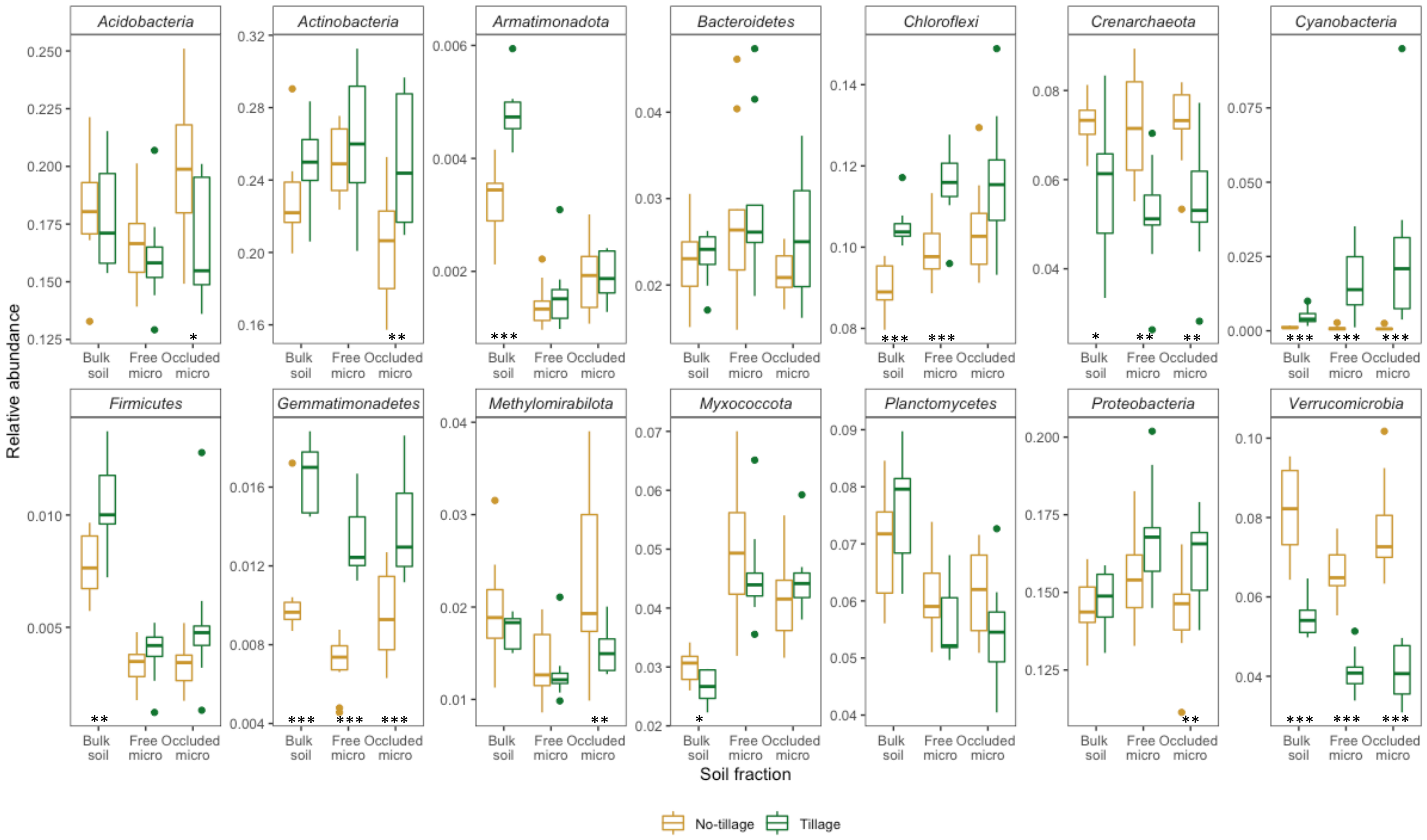

**Figure S11**. Relative abundances of representative phyla at Lancaster, WI. Bulk soil = whole soil; Free micro = microaggregate fraction from bulk soil, 53–250 μm; Occluded micro = microaggregate fraction occluded within macroaggregate fraction, 53–250 μm. Asterisks indicate significant treatment differences in relative abundances between tillage treatment, within fraction, determined using *corncob::differentialTest*, with taxa agglomerated at the phylum level: *** = $p < 0.001$, ** = $p < 0.01$, * = $p < 0.05$.

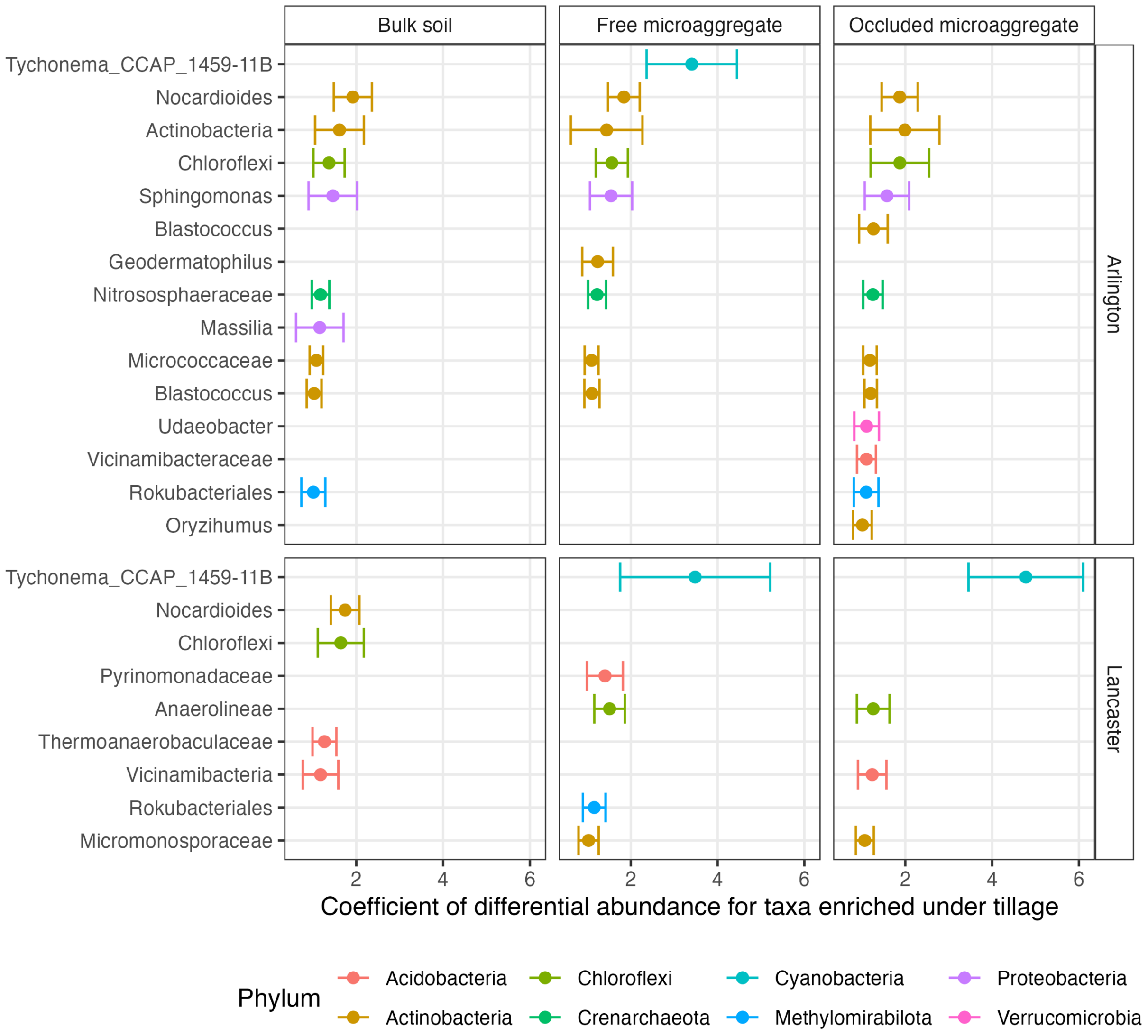

**Figure S12**. Taxa with positive differential abundance (enrichment) under tillage for each soil fraction and site. The x-axis is the coefficient of differential abundance, in this case (showing enrichment), demonstrating an increase in relative abundances of taxa under tillage as compared to no-tillage, with only coefficients > 1.0 plotted. Each point represents a single OTU, colored by phylum, and labelled on the y-axis with the finest available taxonomy; there may be more than one OTU definable to the same name and level of taxonomy. Error bars represent ± 1.96 SE.

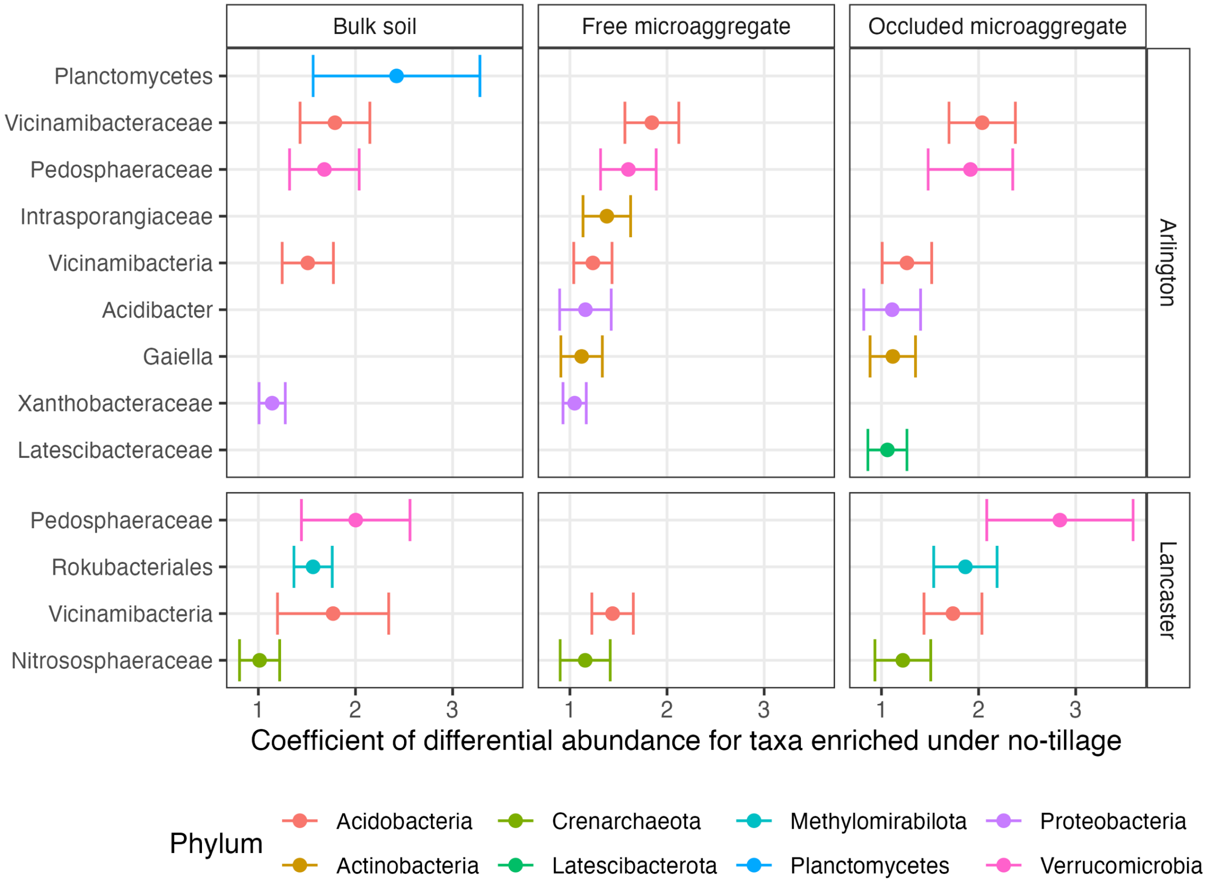

**Figure S13**. Taxa with positive differential abundance (enrichment) under no-tillage for each soil fraction and site. The x-axis is the coefficient of differential abundance, in this case (showing enrichment), demonstrating an increase in relative abundances of taxa under no-tillage as compared to tillage, with only coefficients > 1.0 plotted. Each point represents a single OTU, colored by phylum, and labelled on the y-axis with the finest available taxonomy; there may be more than one OTU definable to the same name and level of taxonomy. Error bars represent ± 1.96 SE.

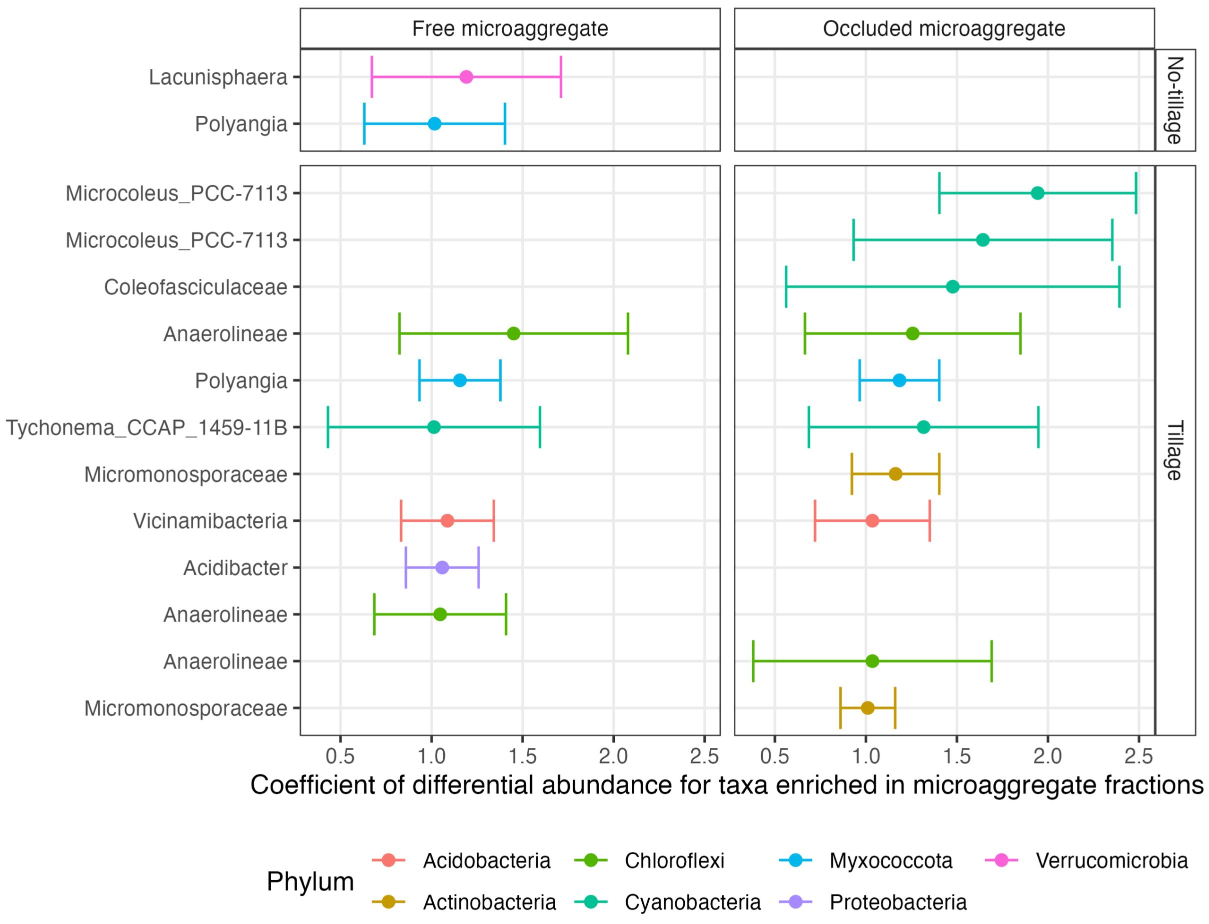

**Figure S14**. Taxa with positive differential abundance (enrichment) in microaggregate fractions at Lancaster, WI. The x-axis is the coefficient of differential abundance, in this case (showing enrichment), demonstrating an increase in relative abundances of taxa in the free and occluded microaggregate fractions, as compared to the bulk soil, with only coefficients > 1.0 plotted. Each point represents a single OTU, colored by phylum, and labelled on the y-axis with the finest available taxonomy; there may be more than one OTU definable to the same name and level of taxonomy, such as *Anaerolineae*. There were no responders that met our criteria at the Arlington site (i.e., the responders were very rare). Error bars represent ± 1.96 SE.

#### SI References

Bouyoucos, G.J., 1962. Hydrometer Method Improved for Making Particle Size Analyses of Soils. Agronomy Journal 54, 464–465. doi:10.2134/agronj1962.00021962005400050028x

Bray, R.H., Kurtz, L.T., 1945. Determination of Total, Organic, and Available Forms of Phosphorus in Soils. Soil Science 59, 39–46. doi:10.1097/00010694-194501000-00006

Campbell, C.D., Chapman, S.J., Cameron, C.M., Davidson, M.S., Potts, J.M., 2003. A Rapid Microtiter Plate Method To Measure Carbon Dioxide Evolved from Carbon Substrate Amendments so as To Determine the Physiological Profiles of Soil Microbial Communities by Using Whole Soil. Applied and Environmental Microbiology 69, 3593–3599. doi:10.1128/aem.69.6.3593-3599.2003

Kozich, J.J., Westcott, S.L., Baxter, N.T., Highlander, S.K., Schloss, P.D., 2013. Development of a Dual-Index Sequencing Strategy and Curation Pipeline for Analyzing Amplicon Sequence Data on the MiSeq Illumina Sequencing Platform. Appl. Environ. Microbiol. 79, 5112–5120. doi:10.1128/aem.01043-13

Richards, L.A., 1954. Diagnosis and Improvement of Saline and Alkali Soils. Soil Science 78, 154. doi:10.1097/00010694-195408000-00012

Schulte, E.E., Hopkins, B.G., 1996. Estimation of Soil Organic Matter by Weight Loss-on-Ignition, in: Magdoff, F.R., Tabatabai, M.A., Hanlon, E.A. (Eds.), Soil Organic Matter: Analysis and Interpretation. Soil Science Society of America, Madison, WI, pp. 21–31. doi:10.2136/sssaspecpub46.c3

Stegen, J.C., Lin, X., Fredrickson, J.K., Chen, X., Kennedy, D.W., Murray, C.J., Rockhold, M.L., Konopka, A., 2013. Quantifying community assembly processes and identifying features that impose them. The ISME Journal 7, 2069–2079. doi:10.1038/ismej.2013.93

Stegen, J.C., Lin, X., Fredrickson, J.K., Konopka, A.E., 2015. Estimating and mapping ecological processes influencing microbial community assembly. Frontiers in Microbiology 6. doi:10.3389/fmicb.2015.00370

Stegen, J.C., Lin, X., Konopka, A.E., Fredrickson, J.K., 2012. Stochastic and deterministic assembly processes in subsurface microbial communities. The ISME Journal 6, 1653–1664. doi:10.1038/ismej.2012.22

Thomas, G.W., 1982. Exchangeable Cations, in: Methods of Soil Analysis, Part 2 Chemical and Microbiological Properties. pp. 159–165. doi:10.2134/agronmonogr9.2.2ed.c9

Walters, W., Hyde, E.R., Berg-Lyons, D., Ackermann, G., Humphrey, G., Parada, A., Gilbert, J.A., Jansson, J.K., Caporaso, J.G., Fuhrman, J.A., Apprill, A., Knight, R., Bik, H., 2016. Improved Bacterial 16S rRNA Gene (V4 and V4-5) and Fungal Internal Transcribed Spacer Marker Gene Primers for Microbial Community Surveys. MSystems 1, e00009-15. doi:10.1128/msystems.00009-15

Willis, A.D., Bunge, J., Whitman, T.L., 2017. Improved detection of changes in species richness in high diversity microbial communities. Journal of the Royal Statistical Society: Series C (Applied Statistics) 66, 963–977. doi:10.1111/rssc.12206
